# A plasma membrane-localized polycystin-1/polycystin-2 complex in endothelial cells elicits vasodilation

**DOI:** 10.1101/2021.10.16.464653

**Authors:** Charles E. Mackay, Miranda Floen, M. Dennis Leo, Raquibul Hasan, Carlos Fernández-Peña, Purnima Singh, Kafait U. Malik, Jonathan H. Jaggar

**Author notes:** Author Contributions: C.M. and J.H.J designed research, C.M., M.F, M.D.M., R.H., C. F-P. and P.S. performed research, C.M., M.F, M.D.M., R.H., C. F-P. and P.S. analyzed data, C.M., K.M. and J.H.J. wrote the paper.

## Abstract

Polycystin-1 (PC-1, PKD1), a receptor-like protein expressed by the *Pkd1* gene, is present in a wide variety of cell types, but its cellular location, signaling mechanisms and physiological functions are poorly understood. Here, by studying tamoxifen-inducible, endothelial cell (EC)-specific *Pkd1* knockout *(Pkd1* ecKO) mice, we show that flow activates PC-1-mediated, Ca^2+^-dependent cation currents in ECs. EC-specific PC-1 knockout attenuates flow-mediated arterial hyperpolarization and vasodilation. PC-1-dependent vasodilation occurs over the entire functional shear stress range and primarily via the activation of nitric oxide synthase (NOS), with a smaller contribution from small-conductance Ca^2+^-activated K^+^ (SK) channels. EC-specific PC-1 knockout increases systemic blood pressure without altering kidney anatomy. PC-1 coimmunoprecipitates with polycystin-2 (PC-2, PKD2), a TRP polycystin channel, and clusters of both proteins locate in nanoscale proximity in the EC plasma membrane. Knockout of either PC-1 or PC-2 *(Pkd2* ecKO mice) abolishes surface clusters of both PC-1 and PC-2 in ECs. Single knockout of PC-1 or PC-2 or double knockout of PC-1 and PC-2 *(Pkd1/Pkd2* ecKO mice) similarly attenuates flow-mediated vasodilation. Flow stimulates non-selective cation currents in ECs that are similarly inhibited by either PC-1 or PC-2 knockout or by interference peptides corresponding to the C-terminus coiled-coil domains present in PC-1 or PC-2. In summary, we show that PC-1 regulates arterial contractility and demonstrate that this occurs through the formation of an interdependent signaling complex with PC-2 in endothelial cells. Flow stimulates PC-1/PC-2 clusters in the EC plasma membrane, leading to Ca^2+^ influx, NOS and SK channel activation, vasodilation and a reduction in blood pressure.

## Introduction

Blood vessels are lined by endothelial cells, which regulate several physiological functions, including contractility, to control regional organ flow and systemic pressure. Endothelial cells can release several diffusible factors, including nitric oxide (NO), which relaxes arterial smooth muscle cells, leading to vasodilation (1). Endothelial cells also electrically couple to smooth muscle cells and directly modulate their membrane potential to modify arterial contractility (2). Several receptor agonists, substances and mechanical stimuli, such as intravascular flow, are known to act in an endothelial cell-dependent manner to regulate arterial functions. In many cases, the molecular mechanisms by which these physiological stimuli activate signaling in endothelial cells to modulate vascular contractility are unclear.

Polycystin-1 (PC-1, PKD1) is a receptor-like transmembrane protein encoded by the *Pkd1* gene (3). PC-1 is expressed in various cell types, including endothelial cells, and is predicted to form eleven transmembrane helices, an extracellular N-terminus and an intracellular C-terminus (3–8). The PC-1 N-terminus is large (>3000 amino acid residues) and contains multiple putative adhesion- and ligand-binding sites (3, 7–11). As such, PC-1 is proposed to act as a mechanical sensor and ligand receptor, although stimuli that activate PC-1 and its functional significance are unclear.

Endothelial cells also express polycystin-2 (PC-2, PKD2), a protein encoded by the *Pkd2* gene (6). PC-2 is a member of the transient receptor potential (TRP) channel family and is also termed TRP polycystin 1 (TRPP1) (12). PC-1 and PC-2 have been proposed to signal through independent and interdependent mechanisms, with much of this knowledge derived from experiments studying recombinant proteins and cultured cells. Supporting their independence, PC-1 and PC-2 exhibit distinct developmental and expression profiles in different kidney cell types (13). PC-2 channels do not require PC-1 to traffic to primary cilia and generate currents in primary-cultured kidney collecting duct cells (14, 15). PC-1 is also proposed to act as an atypical G protein-coupled receptor (16). In addition, C- and N-terminus-deficient PC-2 can alone form a homotetrameric ion channel with each subunit containing six transmembrane domains, when visualized using cryo-EM (17, 18). Evidence supporting PC-1 and PC-2 interdependency includes that mutations in either *Pkd1* or *Pkd2* result in autosomal dominant polycystic kidney disease (ADPKD), the most prevalent monogenic disorder in humans (19). ADPKD is typically characterized by the appearance of renal cysts, but patients can develop hypertension before any kidney dysfunction and cardiovascular disease is the leading (~50 %) cause of death in patients (20–24). Experiments studying recombinant proteins and cultured kidney cell lines have provided evidence that PC-1 and PC-2 can exist in a protein complex (11, 25–31). Several domains in PC-1 and PC-2 may physically interact, including their C-terminus coiled-coils (25, 26, 28, 29, 32). The structure of a PC-1/PC-2 heterotetrameric complex, which forms in a 1:3 stoichiometry, has also been resolved using cryo-EM (25). Despite more than two decades of research, signaling mechanisms and physiological functions of PC-1 and PC-2 and their potential dependency or independence are poorly understood, particularly in extra-renal cell types, such as endothelial cells.

Here, we generated inducible, cell-specific PC-1 knockout *(Pkd1* ecKO) mice to investigate signaling mechanisms and physiological functions of PC-1 in endothelial cells of resistance-size arteries. We also studied endothelial cell-specific PC-2 knockout *(Pkd2* ecKO) mice and produced PC-1/PC-2 double knockout *(Pkd1/Pkd2* ecKO) mice to investigate whether PC-1 acts in an independent manner or is dependent on PC-2 to respond to physiological stimuli and elicit functional responses. Our data demonstrate that PC-1 and PC-2 protein clusters form an interdependent plasma membrane complex in endothelial cells which is activated by flow to produce vasodilation and reduce blood pressure.

## Results

### Generation and validation of tamoxifen-inducible, endothelial cell-specific *Pkd1* knockout mice

Mice with loxP sites flanking exons 2-4 of the *Pkd1* gene *(Pkd1^fl/fl^)* were bred with tamoxifen-inducible endothelial cell-specific Cre mice *(Cdh5-Cre/ERT2)* to generate *Pkd1^fl/fl^:Cdh5-Cre/ERT2* mice. Genomic PCR indicated that tamoxifen (i.p.) stimulated recombination of the *Pkd1* gene in mesenteric arteries of *Pkd1^fl/fl^:Cdh5-Cre/ERT2* mice but did not modify the *Pkd1* gene in arteries of *Pkd1^fl/fl^* controls (Supplementary Fig. 1A). Western blotting demonstrated that tamoxifen treatment reduced PC-1 protein in mesenteric arteries of *Pkd1^fl/fl^:Cdh5-Cre/ERT2* mice to ~66.6 % of that in *Pkd1^fl/fl^* control mice (Fig. 1A-B). This reduction in PC-1 protein in *Pkd1^fl/fl^:Cdh5-Cre/ERT2* mice is expected given that arterial smooth muscle cells also express PC-1 (33). In contrast, tamoxifen did not alter the expression of PC-2, small-conductance Ca^2+^-activated K^+^ (SK3), intermediate-conductance Ca^2+^-activated K^+^ (IK), TRP vanniloid 4 (TRPV4) or Piezo1 channels, endothelial NO synthase (eNOS) or G protein-coupled receptor 68 (GPR68) in arteries of *Pkd1^fl/fl^:Cdh5-Cre/ERT2* mice (Fig. 1A-B). Immunofluorescence imaging of mesenteric arteries en face demonstrated that tamoxifen treatment abolished PC-1 labeling in endothelial cells of *Pkd1^fl/fl^:Cdh5-Cre/ERT2* mice (Fig. 1C). Tamoxifen-treated *Pkd1^fl/fl^:Cdh5-Cre/ERT2* mice will hereafter be referred to as *Pkd1* ecKO mice.

**Figure 1.**
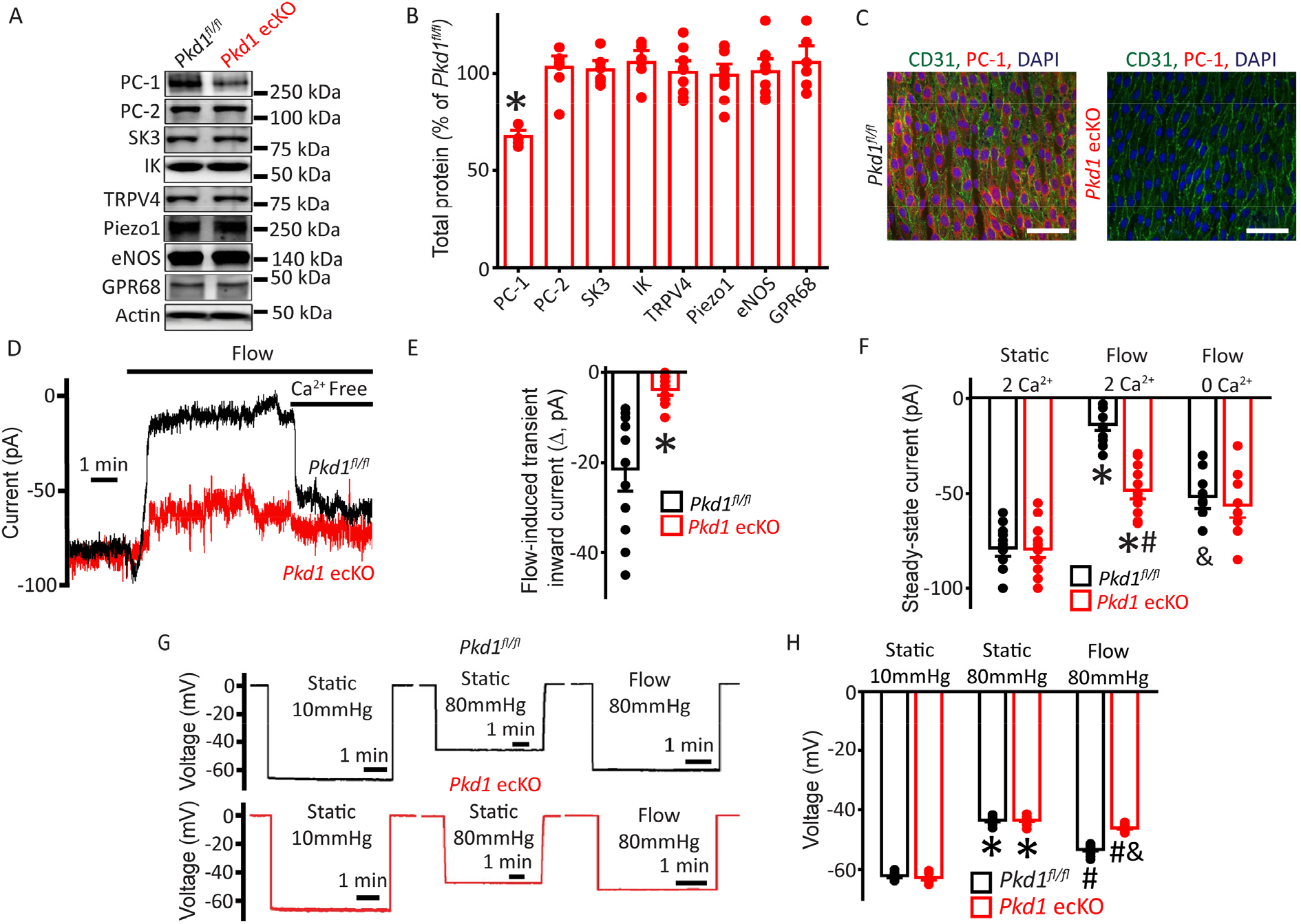
Flow stimulates PC-1-mediated, Ca^2+^-dependent currents in mesenteric artery endothelial cells that elicit arterial hyperpolarization. (A) Representative Western blots illustrating PC-1, PC-2, Piezo1, GPR68, SK3, IK, TRPV4, eNOS and actin proteins in mesenteric arteries of *Pkd1^fl/fl^* and *Pkd1* ecKO mice. (B) Mean data for PC-1, PC-2, Piezo1, GPR68, SK3, IK, TRPV4, eNOS, with n=4, 6, 8, 4, 4, 4, 7, 7, respectively. * indicates P<0.05. (C) En-face immunofluorescence illustrating that PC-1 (Alexa Fluor 546) is abolished in endothelial cells of *Pkd1* ecKO mice mesenteric arteries (representative of 8 arteries from *Pkd1^fl/fl^* and 7 *Pkd1* ecKO mice, respectively). CD31 (Alexa Fluor 488) and DAPI are also shown. Scale bars = 50 μm. (D) Original recordings of steady-state current modulation by flow (10 ml/min) and effect of removing bath Ca^2+^ in endothelial cells of *Pkd1^fl/fl^* and *Pkd1* ecKO mice voltage-clamped at −60 mV. (E) Mean data for flow-induced transient inward current. n=14 for *Pkd1^fl/fl^* and n=16 for *Pkd1* ecKO. * indicates P<0.05 versus *Pkd1^fl/fl^*. (F) Mean data for steadystate currents *(Pkd1^fl/fl^:* static+Ca^2+^, n=15; flow+Ca^2+^, n=15; flow with Ca^2+^ free bath solution, n=12 and *Pkd1* ecKO: static+Ca^2+^, n=16; flow+Ca^2+^, n=16; flow with Ca^2+^ free bath, n=13). *P<0.05 versus static+2 mM Ca^2+^ in the same genotype, # P<0.05 vs *Pkd1^fl/fl^* under the same condition, & P<0.05 versus flow Ca^2+^ in the same genotype. (G) Original membrane potential recordings obtained using microelectrodes in pressurized (80 mmHg) mesenteric arteries of *Pkd1^fl/fl^* and *Pkd1* ecKO mice (H) Mean data *(Pkd1^fl/fl^:* 10 mmHg, n=8; 80 mmHg, n=14; 80 mmHg+flow, n=19; *Pkd1* ecKO: 10 mmHg, n=9; 80 mmHg, n=14; 80 mmHg+flow, n=18). * P<0.05 versus static at 10 mmHg in the same genotype. # P<0.05 for flow versus static at 80 mmHg in the same genotype. & indicates P<0.05 versus *Pkd1^fl/fl^* under the same condition.

### Flow stimulates a PC-1-dependent reduction in inward current in endothelial cells

Patch-clamp electrophysiology was performed to investigate the regulation of plasma membrane currents by PC-1 in mesenteric artery endothelial cells. Currents were recorded at steady-state voltage (−60 mV) using the whole-cell configuration with physiological ionic gradients. In a static bath, endothelial cells of *Pkd1^fl/fl^* mice generated a mean steady-state inward current of ~ −78.9 pA (Fig. 1D, F). Flow stimulated an initial transient increase in inward current of ~-21.4 pA (Fig. 1D-E). This transient increase in inward current was followed by a sustained reduction in inward current that reached steady-state at ~-13.8 pA in *Pkd1^fl/fl^* cells (Fig. 1D, F). During flow, the removal of bath Ca^2+^ increased mean inward current to ~-51.6 pA in *Pkd1^fl/fl^* cells (Fig.1D, F). In a static bath, mean steady-state inward currents were similar in *Pkd1^fl/fl^* and *Pkd1* ecKO cells (Fig. 1D, F). In contrast, the flow-activated transient inward current in *Pkd1* ecKO cells was ~17.9 % of that in *Pkd1^fl/fl^* cells (Fig. 1D-E). Similarly, the sustained flow-mediated reduction in inward current in *Pkd1* ecKO cells was ~47.9% of that in *Pkd1^fl/fl^* cells (Fig. 1D, F). In the continuous presence of flow, the removal of bath Ca^2+^ also resulted in a smaller increase in inward current in *Pkd1* ecKO cells than in *Pkd1^fl/fl^* cells (Fig. 1D, F). Ca^2+^ removal under flow increased inward current only ~ 7.9 pA in *Pkd1* ecKO cells, or ~15.3 % of that in *Pkd1^fl/fl^* cells (Fig. 1D, F). These data demonstrate that flow stimulates a PC-1-dependent biphasic response composed of an initial transient inward current followed by a sustained Camdependent reduction in inward current in endothelial cells.

### Endothelial cell PC-1 contributes to flow-mediated arterial hyperpolarization

The flow-mediated, PC-1-dependent reduction in inward current in endothelial cells suggested that PC-1 may regulate arterial membrane potential, a major determinant of contractility (34). Arterial membrane potential was measured by impaling sharp glass microelectrodes into pressurized (80mmHg) third-order mesenteric arteries of *Pkd1^fl/fl^* and *Pkd1* ecKO mice. In the absence of intraluminal flow, the membrane potentials of *Pkd1^fl/fl^* and *Pkd1* ecKO arteries were similar at either low (10 mmHg) or physiological (80 mmHg) pressures (Fig. 1G, H). At 80 mmHg, intraluminal flow hyperpolarized *Pkd1^fl/fl^* arteries by ~10 mV, but *Pkd1* ecKO arteries by only ~ 2.6 mV, or ~ 27.1 % of that in controls (Fig. 1G, H). These data suggest that PC-1 in endothelial cells contributes to flow-mediated arterial hyperpolarization.

### PC-1 activates eNOS and SK channels in endothelial cells to elicit vasodilation

The regulation of contractility by endothelial cell PC-1 was measured in pressurized (80 mmHg) third-order myogenic mesenteric arteries. Intravascular flow (15 dyn/cm^2^) stimulated sustained and fully reversible dilations in mesenteric arteries of both *Pkd1^fl/fl^* and *Pkd1* ecKO mice (Fig. 2A). However, in *Pkd1* ecKO arteries, dilations to on-off flow stimuli were ~34.8 % of those in *Pkd1^fl/fl^* arteries (Fig. 2A, B). In contrast, ACh-induced vasodilation was similar in *Pkd1^fl/fl^* and *Pkd1* ecKO arteries (Fig. 2B, Supplementary Fig. 2A). To determine the functional flow range for PC-1 in endothelial cells, we measured vasoregulation to shear stresses between 3 and 35 dyn/cm^2^ (35). Cumulative stepwise increases in flow caused progressive dilation, with a maximum at 27 dyn/cm^2^ in *Pkd1^fl/fl^* arteries (Fig. 2C, D). Increasing shear stress above 27 dyn/cm^2^ slightly attenuated maximal vasodilation (Fig. 2C, D). The PC-1 sensitive component of flow-mediated dilation was calculated by subtracting responses in *Pkd1* ecKO arteries from those in *Pkd1^fl/fl^* arteries. Flow-mediated dilation in *Pkd1* ecKO arteries was attenuated over the entire shear-stress range to between ~38.0 and 50.0 % of that in *Pkd1^fl/fl^* arteries (Fig. 2C, D, Supplementary Fig. 2B). Myogenic tone, depolarization (60mM K^+^)-induced constriction and passive diameter were similar in *Pkd1^fl/fl^* and *Pkd1* ecKO arteries (Supplementary Fig. 2C-F). These data indicate that a broad intravascular flow range activates PC-1 in endothelial cells to induce vasodilation.

**Figure 2.**
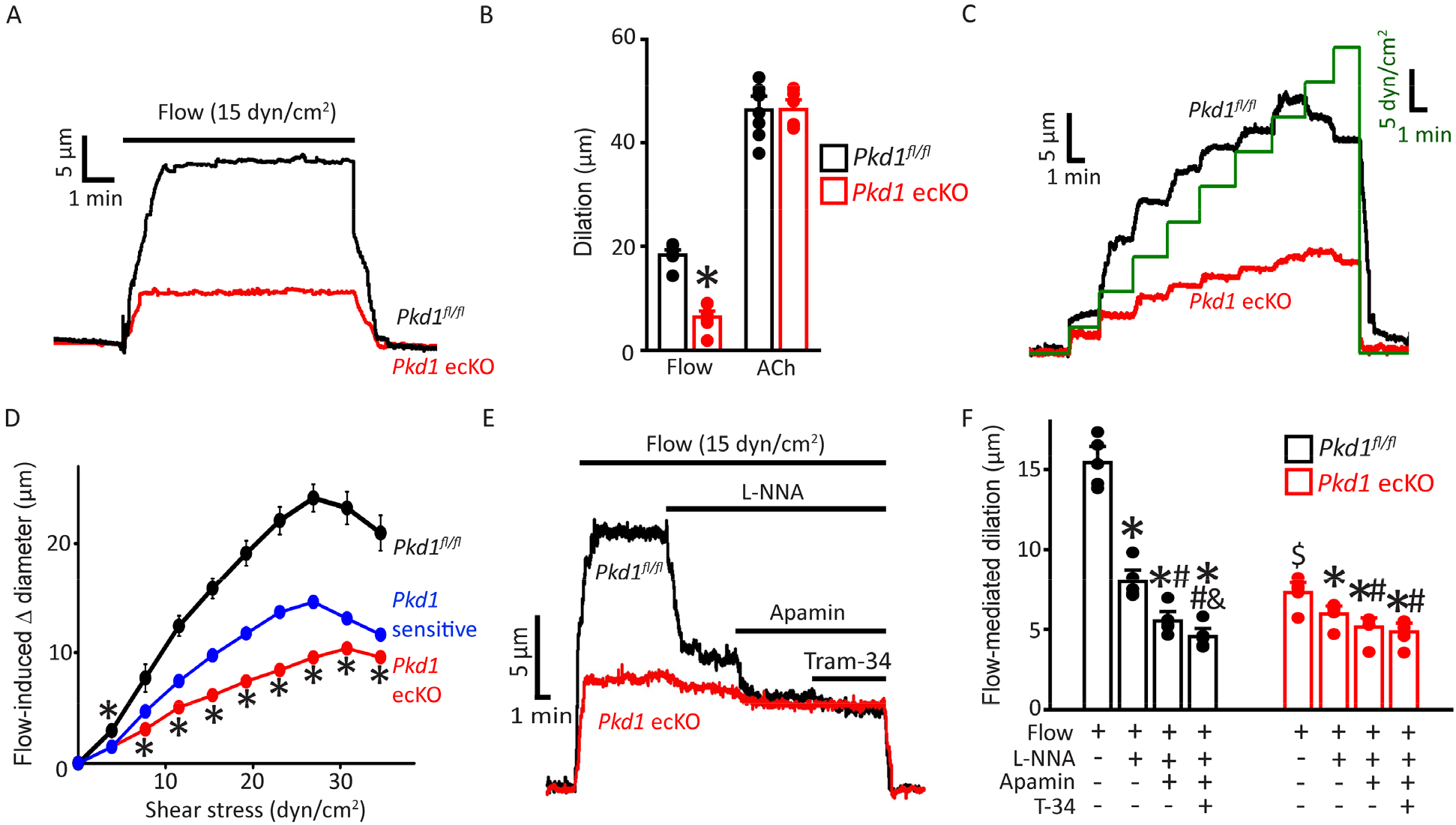
Endothelial cell PC-1 stimulates vasodilation primarily via NOS activation. (A) Representative traces illustrating reversible flow-mediated dilation in pressurized (80 mmHg) mesenteric arteries of *Pkd1^fl/fl^* and *Pkd1* ecKO mice. (B) Mean dilation to flow (15 dyn/cm^2^) or ACh (10 μM). *P<0.05 versus *Pkd1^fl/fl^*. n=8 for each dataset. (C) Representative diameter changes to stepwise increases in intravascular flow in pressurized (80 mmHg) mesenteric arteries from *Pkd1^fl/fl^* and *Pkd1* ecKO mice. (D) Mean data. The *Pkd1* -sensitive component of flow-mediated vasodilation is shown in blue. n=4 each for *Pkd1^fl/fl^* and *Pkd1* ecKO. * P<0.05 versus *Pkd1^fl/fl^.* (E) Regulation of flow (15 dyn/cm^2^)-mediated dilation by L-NNA (10 μM), apamin (300 nM) and Tram-34 (300 nM) in pressurized (80 mmHg) mesenteric arteries of *Pkd1^fl/fl^* and *Pkd1* ecKO mice. (F) Mean data. n=5 for each dataset. Symbols illustrate P<0.05 for: $ versus flow in *Pkd1fl/fl*, * versus flow in the same genotype. # versus flow+L-NNA in the same genotype. & versus flow+L-NNA+apamin in the same genotype.

Ca^2+^ influx activates NOS, SK and IK channels in endothelial cells, leading to vasodilation (1,2). Next, we studied contributions of each of these proteins to flow-mediated, PC-1-dependent vasodilation. L-NNA, a NOS inhibitor, reduced mean flow-mediated dilation to ~48.2% of control in pressurized (80 mmHg) myogenic *Pkd1^fl/fl^* arteries (Fig. 2E, F). Coapplication of apamin, a SK channel blocker, further reduced flow-mediated dilation to ~35.8% of control. The addition of Tram-34, an IK inhibitor, only slightly reduced flow-mediated vasodilation to ~29.5% of controls (Fig. 2E, F). PC-1 knockout reduced the L-NNA- and apamin-sensitive components of flow-mediated dilation to ~37.6% and ~50.8% of those in *Pkd1^fl/fl^* arteries (Fig. 2E, F). In contrast, the Tram-34-sensitive component of flow-mediated dilation was not altered by PC-1 knockout (Fig. 2E, F). We have previously shown that in the absence of intravascular flow, L-NNA, apamin and Tram-34 do not alter the diameter of pressurized mesenteric arteries (6). These data indicate that flow stimulates PC-1 in endothelial cells, leading to NOS and SK channel activation, which produces vasodilation.

### Endothelial cell PC-1 reduces systemic blood pressure

Radiotelemetry measurements were performed to measure blood pressure in freely-moving, conscious mice. Mean arterial pressure (MAP) was ~93.5 mmHg in *Pkd1^fl/fl^* mice and ~109.1 mmHg in *Pkd1* ecKO mice, or 16.6 % higher (Fig. 3A-B). Underlying this increase in MAP were higher systolic and diastolic blood pressures in *Pkd1* ecKO mice (Supplemental Fig. 3A). In contrast, heart rate and activity were similar in *Pkd1^fl/fl^* and *Pkd1* ecKO mice, as were proximal tubule diameter and glomerular surface area measured in H&E-stained kidney sections (Fig. 3C-F, Supplementary Fig. 3B). These results suggest that flow stimulates PC-1 in endothelial cells, leading to vasodilation and a reduction in systemic blood pressure.

**Figure 3.**
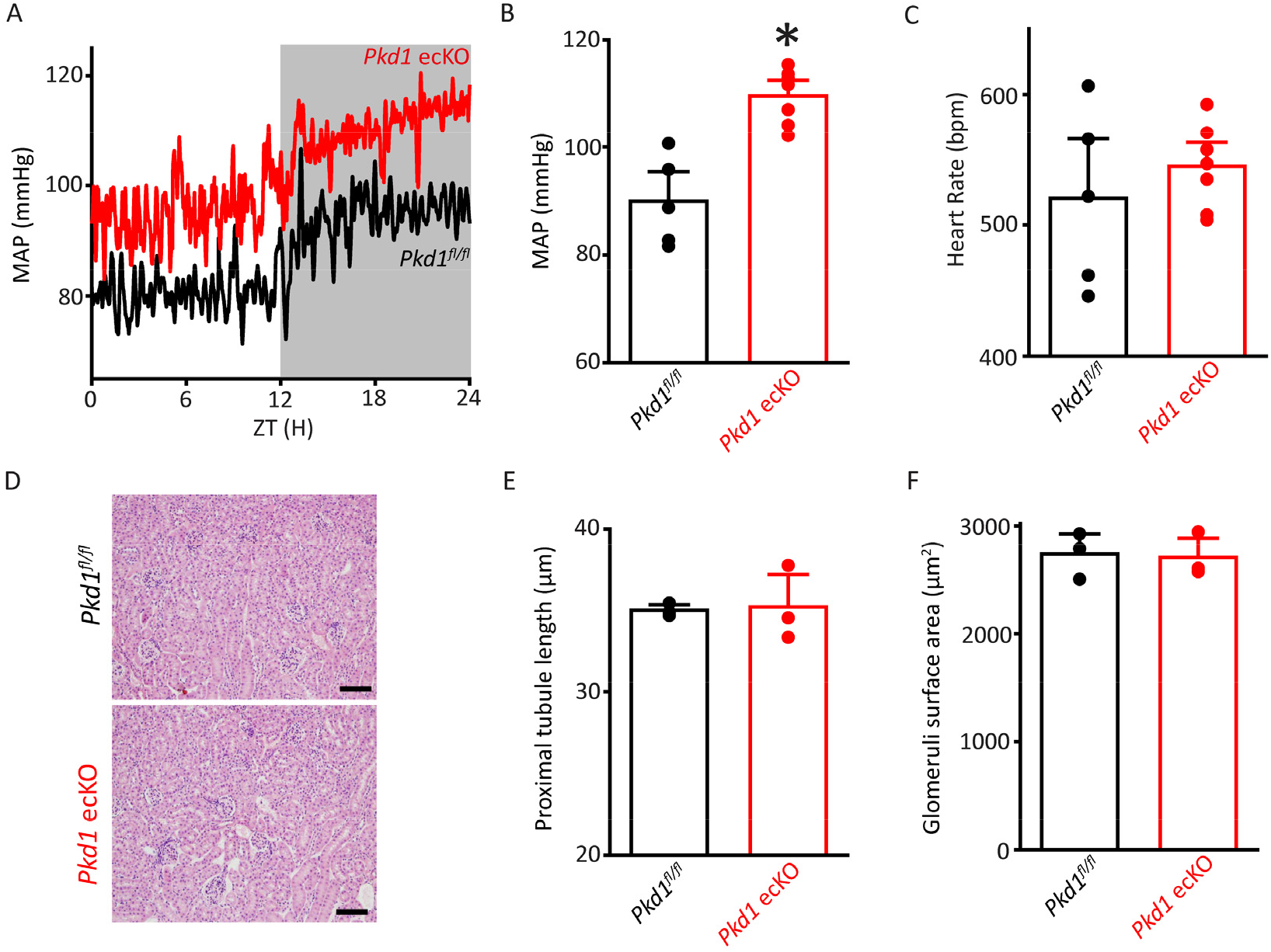
*Pkd1* ecKO mice are hypertensive with normal kidney anatomy. (A) Blood pressure recordings obtained over 24 hours in a *Pkd1^fl/fl^* and *Pkd1* ecKO mouse. (B) Mean arterial pressures (MAP) in *Pkd1^fl/fl^* (n=5) and *Pkd1* ecKO (n=7) mice. * P<0.05 versus *Pkd1^fl/fl^.* (C) Mean heart rate (HR). *Pkd1^fl/fl^*, n=5, *Pkd1* ecKO, n=7. (D) Images of H&E-stained kidney cortex used for histological measurements. Scale bars = 100 μm (F) Mean proximal tubule length. n=15 proximal tubules measured in each mouse, n=3 mice. (G) Mean glomeruli surface area. n=75 glomeruli measured from each mouse, n=3 mice.

### PC-1 and PC-2 coassemble and colocalize in endothelial cells

The vascular phenotype we describe here for *Pkd1* ecKO mice is similar to that of *Pkd2* ecKO mice (6). Thus, we tested the hypothesis that PC-1 and PC-2 are components of the same flow-sensitive signaling pathway in endothelial cells. PC-1 and PC-2 coimmunoprecipitated in mesenteric artery lysate, suggesting they coassemble (Fig. 4A). Next, we used several different imaging techniques to measure the spatial proximity of PC-1 and PC-2 proteins in endothelial cells. Immunofluorescence energy transfer (immunoFRET) microscopy using Alexa Fluor 546 and Alexa Fluor 488-tagged secondary antibodies bound to PC-1 and PC-2 primary antibodies, respectively, generated mean N-FRET of 28.4±1.7% in endothelial cells (n=12, P<0.05, Fig. 4B). The Förster coefficient of the Alexa Fluor pair used is ~6.3 nm, suggesting PC-1 and PC-2 locate in close spatial proximity. Lattice Structured Illumination super-resolution Microscopy (Lattice-SIM) was used to measure colocalization of PC-1 and PC-2 in endothelial cells of en face mesenteric arteries and in isolated mesenteric artery endothelial cells (Fig. 4C-F). Endothelial cells were identified through immunolabeling of CD31, an endothelial cell-specific marker (Fig. 4C, E). Analysis of these data using both Pearson’s and Mander’s coefficients also indicated that PC-1 and PC-2 colocalize in endothelial cells (Fig. 4D, F) (36).

**Figure 4.**
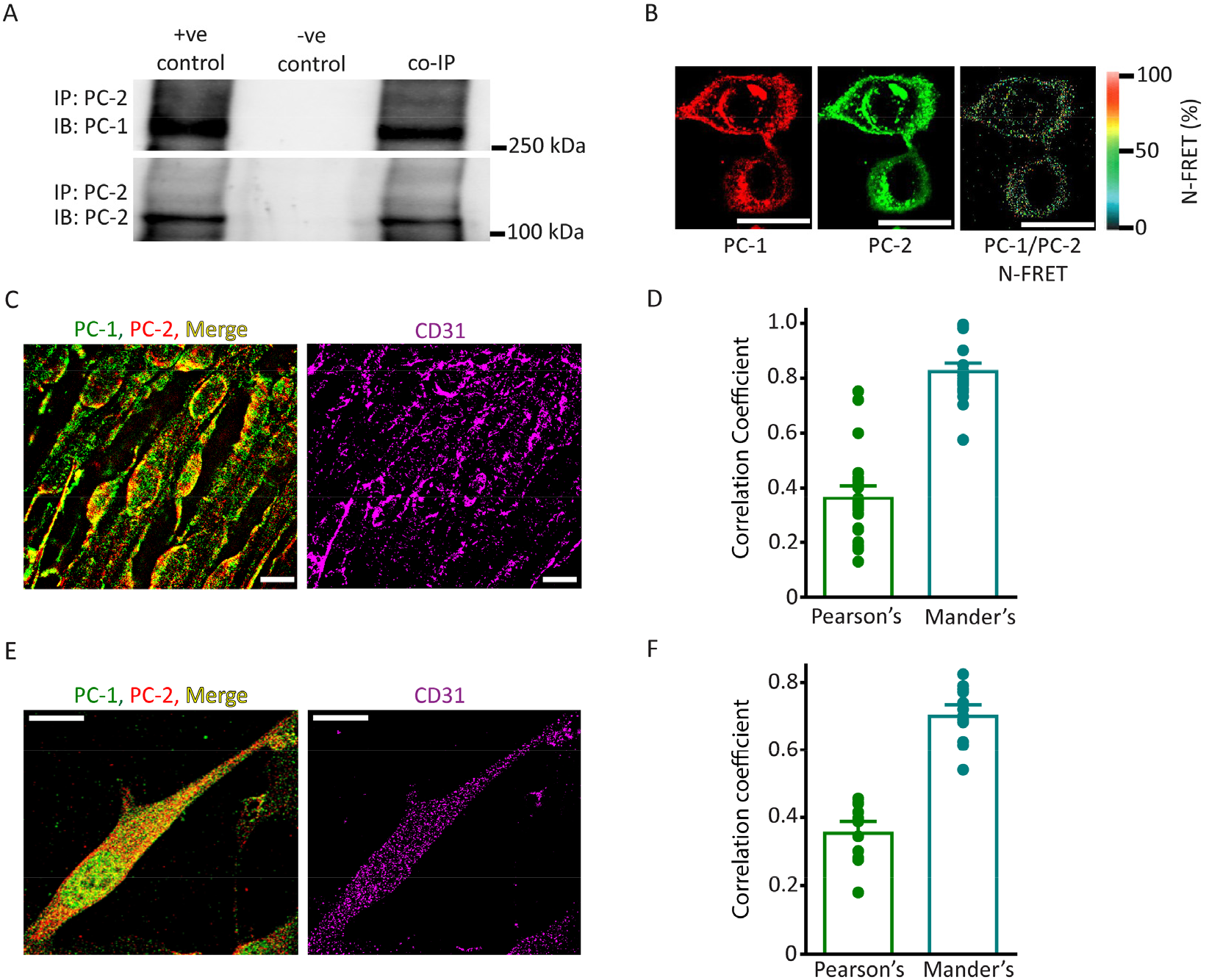
PC-1 and PC-2 coassemble and colocalize in endothelial cells. (A) Representative Western blots illustrating the detection (IB) of both PC-1 and PC-2 (IB) in PC-2 immunoprecipitate (IP) (n=5). (B). PC-1 (Alexa546) and PC-2 (Alexa488) immunofluorescence generate FRET in mesenteric artery endothelial cells. Scale bars = 10 μm. (C) Lattice SIM images of PC-1, PC-2 and CD31 immunofluorescence in the same endothelial cells of an en face mesenteric artery. The merged image is also shown with yellow pixels illustrating colocalization of PC-1 and PC-2. Scale bars = 10 μm. (D) Mean data for PC-1 and PC-2 colocalization using both Pearson’s and Mander’s correlation coefficients. n=25 images, 12 arteries and 6 mice for each dataset. (E) Lattice SIM image of PC-1 and PC-2 immunofluorescence in a mesenteric artery endothelial cell. Yellow pixels illustrate PC-1 to PC-2 colocalization. Scale bar = 10 μm. (F) Mean data for PC-1 and PC-2 colocalization when using Pearson’s and Mander’s correlation coefficients. n=13, 4 mice for each dataset.

### PC-1 and PC-2 surface clusters colocalize and are interdependent in endothelial cells

Single-Molecule Localization Microscopy (SMLM) in combination with Total Internal Reflection Fluorescence (TIRF) imaging was used to measure the properties and nanoscale proximity of PC-1 and PC-2 protein clusters in the plasma membrane of endothelial cells. Localization precision of the Alexa Fluor 488 and Alexa Fluor 647 fluorophores conjugated to secondary antibodies were 29.6±0.6 (n=53) and 24.6±0.7 (n=45) nm, respectively, when imaged in endothelial cells (Supplementary Fig. 4A, B). Discrete clusters of PC-1 and PC-2 were observed in the plasma membrane of *Pkd1^fl/fl^* endothelial cells (Fig. 5A). PC-1 and PC-2 clusters exhibited similar densities (Fig. 5B). The sizes (area) of individual PC-1 and PC-2 clusters were exponentially distributed, with means of ~399.6 and 243.9 nm^2^, respectively (Fig. 5C, Supplementary Fig. 5A, B). Histograms were constructed that contained the distance between the center of each PC-1 cluster and that of its nearest PC-2 neighbor. These data were exponentially distributed (Supplementary Fig. 5C). The mean PC-1 to PC-2 intercentroid distance was ~41.7 nm in *Pkd1^fl/fl^* cells (Fig. 5D). Approximately 29.0 % of PC-1 clusters overlapped with a PC-2 cluster and ~34.8 % of PC-2 clusters overlapped with a PC-1 cluster in *Pkd1^fl/fl^* endothelial cells (Fig. 5E, F, Supplementary Fig. 5D). When experimental data were randomized using Costes’ simulation algorithm (36), PC-1 to PC-2 and PC-2 to PC-1 overlap were only ~9.2 and 9.6 %, respectively in *Pkd1^fl/fl^* cells (Fig. 5E, F, Supplementary Fig. 5D). Flow did not alter PC-1 or PC-2 cluster density, PC-1 or PC-2 cluster size, the distance between PC-1 and PC-2 centroids, or their overlap (Fig. 5B-F, Supplementary Fig. 5D, E). In *Pkd1* ecKO cells, the density of PC-1 and PC-2 clusters was far lower, at ~ 14.0 and 25.0 %, respectively of those in endothelial cells of *Pkd1^fl/fl^* mice (Fig. 5A, B). PC-1 and PC-2 clusters were also smaller and the PC-1 to PC-2 and PC-2 to PC-1 distances were far greater in *Pkd1* ecKO than *Pkd1^fl/fl^* cells (Fig. 5C, D, Supplementary Fig. 5B-D). Furthermore, only ~4.9 % of PC-1 clusters overlapped with a PC-2 cluster in *Pkd1* ecKO cells, with similar results for PC-2 to PC-1 overlap (Fig. 5E, F, Supplementary Fig. 5D). Costes’ randomization of data did not alter PC-1 to PC-2 or PC-2 to PC-1 overlap in *Pkd1* ecKO cells. These data indicate that: 1) PC-1 and PC-2 surface protein clusters are colocalized in endothelial cells, 2) PC-1 or PC-2 knockout inhibits surface expression of both PC-1 and PC-2 proteins and 3) fluorescent clusters in *Pkd1* ecKO cells are likely due to non-specific labeling by secondary antibodies.

**Figure 5.**
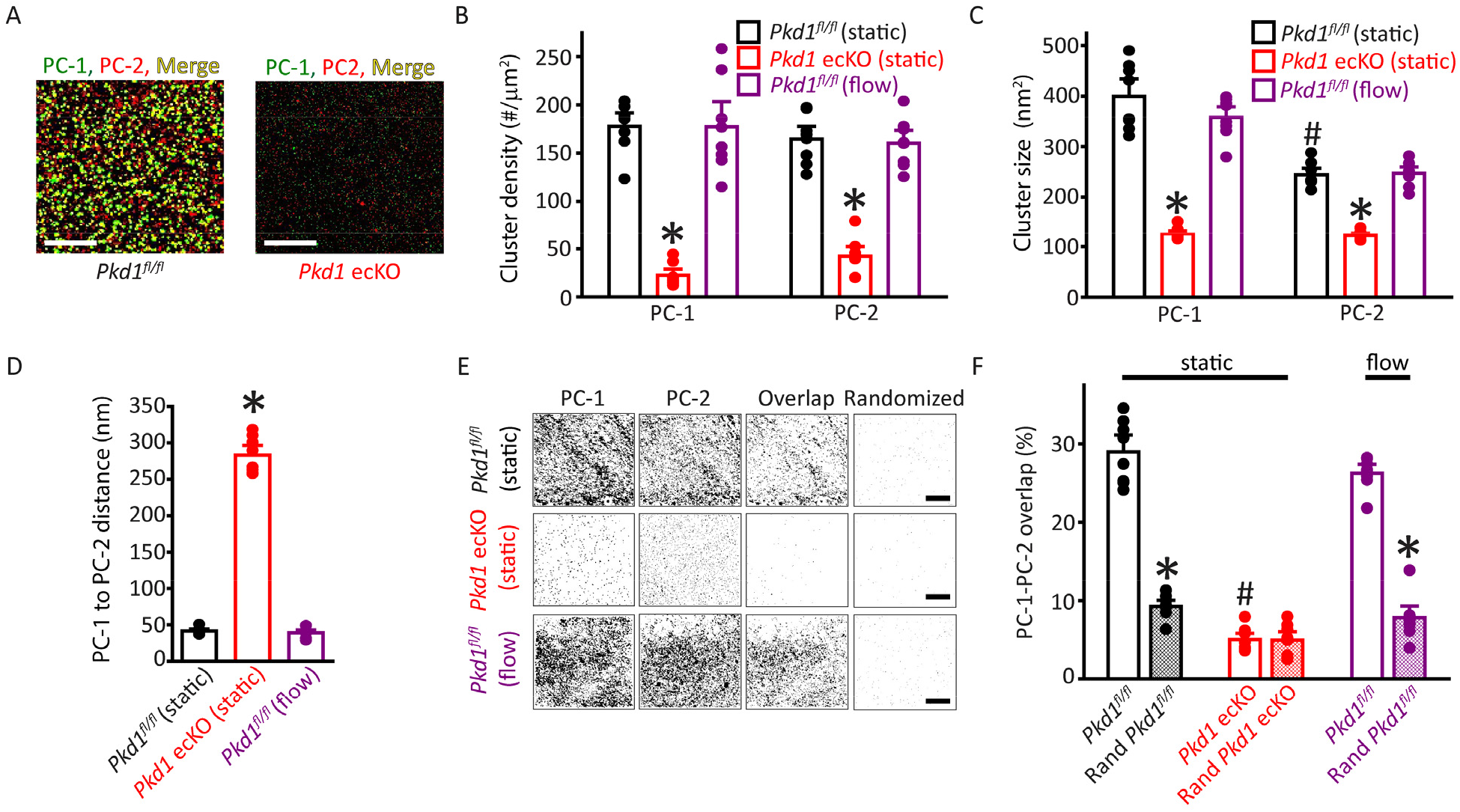
Plasma membrane PC-1 and PC-2 clusters colocalize in endothelial cells. (A) TIRF-SMLM images of PC-1 and PC-2 surface clusters in a *Pkd1^fl/fl^* and *Pkd1* ecKO endothelial cell. Scale bars = 5 μm. (B) Mean data for PC-1 and PC-2 cluster density measured in *Pkd1^fl/fl^* and *Pkd1* ecKO endothelial cells under static and flow (10 ml/min) conditions. n=8 for each dataset. * P<0.05 versus *Pkd1^fl/fl^* (static). (C) Mean data for PC-1 and PC-2 cluster sizes measured in *Pkd1^fl/fl^* and *Pkd1* ecKO endothelial cells under static and flow (10 ml/min) conditions. n=8 for each dataset. * P<0.05 versus respective floxed control in static condition, # P<0.05 versus *Pkd1^fl/fl^* static. (D) Mean data for PC-1 to PC-2 nearest-neighbor analysis. n=8 for each dataset. * P<0.05 versus *Pkd1^fl/fl^* static and *Pkd1^fl/fl^* flow. (E) TIRF-SMLM images of PC-1 and PC-2 clusters, overlap of PC-1 and PC-2 data and overlap of PC-1 and PC-2 data following Coste’s randomization simulation in *Pkd1^fl/fl^* and *Pkd1* ecKO cells. Scale bars = 5 μm. (F) Mean experimental and Costes’ randomized (Rand) data for PC-1 to PC-2 overlap in *Pkd1^fl/fl^* cells in static and flow and *Pkd1* ecKO cells in static. n=8 for each dataset. * P<0.05 versus respective floxed control in static condition, # P<0.05 versus *Pkd1^fl/fl^* static.

To determine whether data obtained using SMLM were specific to endothelial cells of *Pkd1^fl/fl^* and *Pkd1* ecKO mice, we performed similar experiments using tamoxifen-inducible, endothelial cell-specific PC-2 knockout (*Pkd2* ecKO) mice and their controls (*Pkd2^fl/fl^*). The mean densities and sizes of PC-1 and PC-2 clusters, PC-1 to PC-2 and PC-2 to PC-1 intercluster distances and PC-1 to PC-2 and PC-2 to PC-1 overlap were similar in endothelial cells of *Pkd2^fl/fl^* mice and *Pkd1^fl/fl^* mice (Supplementary Fig. 6A-F and Supplementary Fig. 7A, B). Similarly to observations in *Pkd1^fl/fl^* cells, flow did not alter the densities, sizes, intercluster distances or overlap of PC-1 or PC-2 clusters in *Pkd2^fl/fl^* cells (Supplementary Figs. 6A-F and 7A, B). In *Pkd2* ecKO cells, PC-1 and PC-2 cluster densities were far lower, at ~16.2 and 12.6 % of those in *Pkd2^fl/fl^* cells (Supplementary Fig. 6A, B). PC-1 and PC-2 clusters were smaller and the mean distance from PC-2 clusters to their nearest PC-1 neighbor was far greater in *Pkd2* ecKO cells than in *Pkd2^fl/fl^* cells (Supplementary Fig. 6C-F). PC-1 to PC-2 and PC-2 to PC-1 overlap were also lower in *Pkd2* ecKO than *Pkd2^fl/fl^* cells (Supplementary Fig. 7A, B). These data demonstrate that PC-1 and PC-2 surface clusters colocalize in endothelial cells. Knockout of either PC-1 or PC-2 inhibits the surface expression of both PC-1 and PC-2 proteins, supporting their interdependency. Fluorescent clusters observed in *Pkd1* ecKO and *Pkd2* ecKO cells appear to represent non-specific secondary antibody labeling.

### Flow-mediated vasodilation is similarly attenuated in *Pkd1* ecKO and *Pkd1/Pkd2* ecKO arteries

Next, we investigated the functional interdependency of PC-1 and PC-2 in flow-mediated vasodilation. *Pkd1^fl/fl^:Cdh5-Cre/ERT2* and *Pkd2^fl/fl^:Cdh5-Cre/ERT2* mice were crossed to generate a *Pkd1^fl/fl^/Pkd2^fl/fl^:Cdh5-Cre/ERT2* line. Control Cre-negative *Pkd1^fl/fl^/Pkd2^fl/fl^* mice were generated using a similar breeding strategy. Tamoxifen-treatment reduced PC-1 and PC-2 proteins in mesenteric arteries of *Pkd1^fl/fl^/Pkd2^fl/fl^:Cdh5-Cre/ERT2* mice to ~60.8±10.1 and 66.0±9.6 %, respectively of those in *Pkd1^fl/fl^/Pkd2^fl/fl^* mice (Fig. 6A, B, P<0.05 and n=5 for each). In contrast, eNOS was similar in *Pkd1/Pkd2* ecKO arteries, at 111.9±18.5 % of that in *Pkd1^fl/fl^/Pkd2^fl/fl^* (Fig. 6A, B, P>0.05 and n=5 for each). Intravascular flow stimulated vasodilations in pressurized *Pkd1^fl/fl^/Pkd2^fl/fl^* arteries that were similar in magnitude to those in arteries of both *Pkd1^fl/fl^* and *Pkd2^fl/fl^* mice (Figs. 2A-C, 6C, D and ref. (6)). Mean flow-mediated vasodilation in *Pkd1/Pkd2* ecKO mouse arteries was ~39.2-50.3 % of that in *Pkd1^fl/fl^/Pkd2^fl/fl^* arteries (Fig. 6C, D). This inhibition of flow-mediated dilation is similar to that in arteries of *Pkd1* ecKO and *Pkd2* ecKO mice when compared to their respective controls (Figs. 2A-C, 6C, D and ref. (6)). In contrast, myogenic tone, 60 mM K^+^-induced constriction, dilations to ACh or sodium nitroprusside (SNP), a nitric oxide donor, and passive diameter were similar in *Pkd1^fl/fl^/Pkd2^fl/fl^* and *Pkd1/Pkd2* ecKO arteries (Fig. 6C, Supplementary Fig. 8A-E). Collectively, these data indicate that intravascular flow stimulates vasodilation via a mechanism that involves both PC-1 and PC-2 in endothelial cells.

**Fig. 6.**
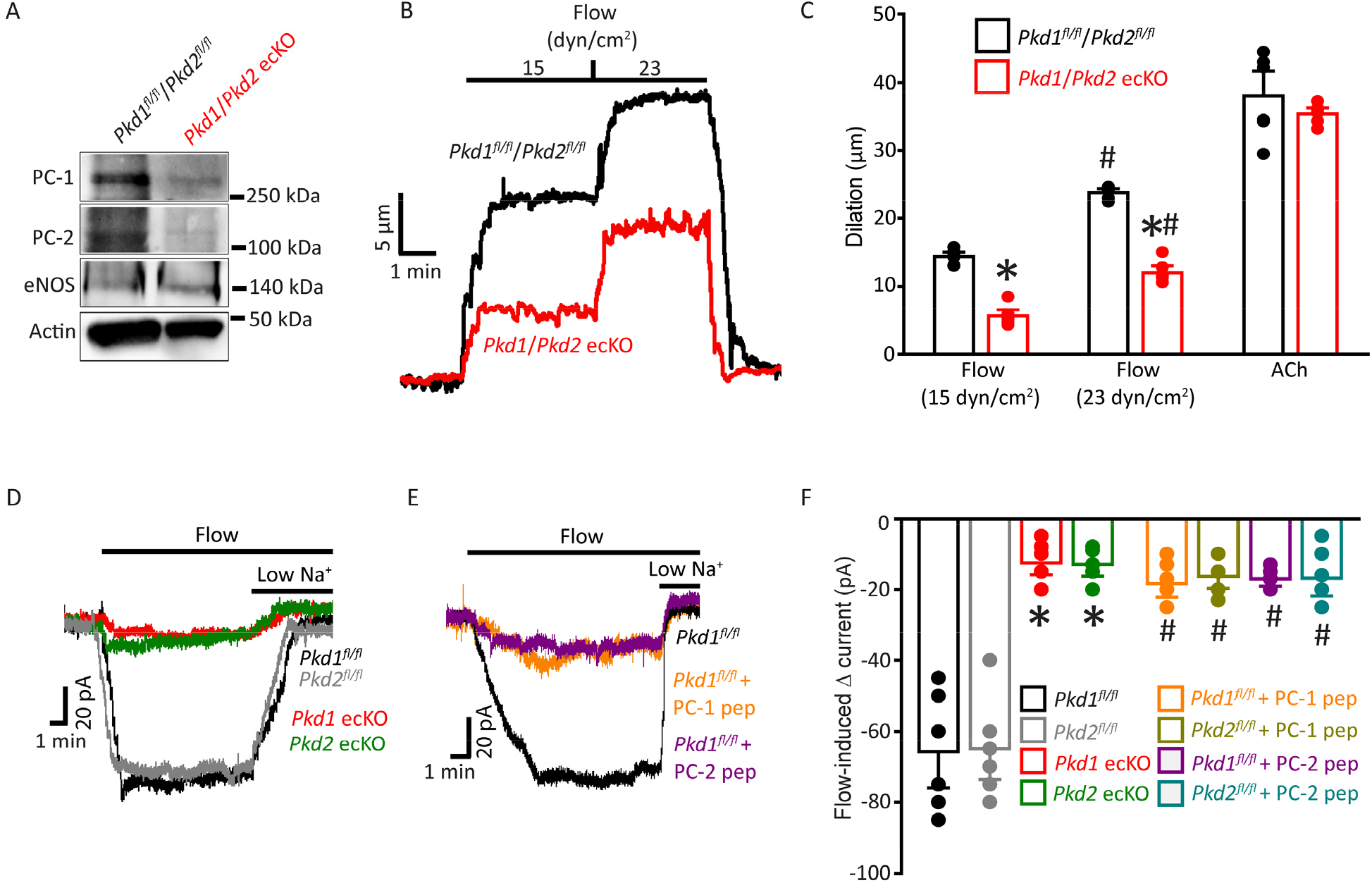
PC-1 and PC-2 are interdependent for flow-mediated vasodilation and I_Cat_ activation in mesenteric artery ECs. (A) Representative Western blots of PC-1, PC-2 and eNOS in mesenteric arteries of *Pkd1^fl/fl^/Pkd2^fl/fl^* and *Pkd1/Pkd2* ecKO mice. (B) Representative traces illustrating flow-mediated dilation by 15 and 23 dyn/cm^2^ in pressurized (80 mmHg) mesenteric arteries from *Pkd1^fl/fl^/Pkd2^fl/fl^* and *Pkd1/Pkd2* ecKO mice. (C) Mean dilation to flow (15 an^d 23^ dyn/cm^2^) or ACh (10 μM). n=6 each for 15 dyn/cm^2^, 23 dyn/cm^2^ and ACh. *P<0.05 versus *Pkd1^fl/fl^/Pkd2^fl/fl^.* #P<0.05 versus 15 dyn/cm^2^ in the same genotype. (D) Original recordings illustrating that flow (10 ml/min) activates I_Cat_s in *Pkd1^fl/fl^* and *Pkd2^fl/fl^* endothelial cells that are similarly reduced in PC-1 ecKO and PC-2 ecKO endothelial cells. (E) Intracellular introduction via the patch pipette of either a PC-1 or PC-2 C-terminus coiled-coil domain peptide reduces flow-induced I_Cat_s in *Pkd1^fl/fl^* and *Pkd2^fl/fl^* endothelial cells. (F) Mean data. PC-1 pep and PC-2 pep indicate peptides corresponding to the coiled-coil domain in PC-1 or PC-2, respectively. *Pkd1^fl/fl^* n=6, *Pkd2^fl/fl^* n=6, *Pkd1* ecKO n=7, *Pkd2* ecKO n=5, *Pkd1^fl/fl^+PC-1* peptide n=6, *Pkd2^fl/fl^+PC-1* peptide n=6, *Pkd1^fl/fl^+PC-2* peptide n=5 and *Pkd2^fl/fl^+PC-2* peptide n=6. * P<0.05 versus respective flox control genotype with no peptide used. # P<0.05 versus control (no peptide) in same genotype.

### PC-1 knockout, PC-2 knockout and coiled-coil domain peptides similarly inhibit flow-activated non-selective cation current (I_Cat_) in endothelial cells

Recombinant PC-1 and PC-2 can physically interact via several domains, including their C-terminal coiled-coils (26, 28, 29, 32). To investigate the functional significance of the PC-1 and PC-2 coiled-coil domains to flow-mediated signaling, we measured current regulation in endothelial cells. To reduce the contribution of other types of ion channels to currents, I_Cat_ was measured using solutions that inhibit K^+^ channels and a bath solution that was Ca^2+^-free. At a steady-state holding potential of −60 mV, flow reversibly stimulated I_Cat_s of similar amplitudes in *Pkd1^fl/fl^* and *Pkd2^fl/fl^* endothelial cells (Fig. 6D, F). In *Pkd1* ecKO endothelial cells, mean flow-activated I_Cat_ was ~19.1 % of that in *Pkd1^fl/fl^* controls (Fig. 6D, F). Flow-activated I_Cat_ in *Pkd2* ecKO cells was similarly reduced, at ~20.0 % of that in *Pkd2^fl/fl^* cells (Fig. 6D, F). Peptides were constructed that correspond to regions within the C-terminal coiled-coil domains in PC-1 (amino acids 4216-4233) and PC-2 (amino acids 874-883) that physically interact (26, 28–30). Intracellular introduction via the pipette solution of either the PC-1 or PC-2 coiled-coil domain peptide similarly reduced flow-activated I_Cat_s to between ~25.8 and 27.8 % of control currents in both *Pkd1^fl/fl^* and *Pkd2^fl/fl^* endothelial cells (Fig. 6E, F). Lowering bath Na^+^ concentration inhibited flow-activated currents in all genotypes and under all conditions. These data indicate that PC-1 and PC-2 are interdependent for flow to activate I_Cat_ in endothelial cells. Data also suggest that physical coupling of PC-1 and PC-2 C-termini is necessary for flow to activate I_Cat_.

## Discussion

Here, we investigated the regulation of arterial contractility by PC-1 and the potential involvement of PC-2 in mediating responses in endothelial cells. Our data show that flow stimulates PC-1-dependent cation currents in endothelial cells that induce arterial hyperpolarization, vasodilation and a reduction in blood pressure. PC-1-dependent vasodilation occurs over the entire functional shear stress range, primarily due to NOS activation, with a smaller contribution through the stimulation of SK channels. PC-1 and PC-2 proteins coassemble and their surface clusters colocalize. Knockout of either PC-1 or PC-2 abolishes surface expression of both proteins. Flow activates a PC-1- and PC2-dependent I_Cat_ that is inhibited by peptides corresponding to the C-terminus coiled-coil domains on either polycystin protein. These data demonstrate that PC-1 regulates arterial contractility by forming an interdependent complex with PC-2 in endothelial cells. Flow stimulates plasma membrane-localized PC-1/PC-2 clusters, leading to Ca^2+^ influx, NOS and SK channel activation, vasodilation and a reduction in blood pressure.

Intraluminal flow stimulates endothelial cells to induce vasodilation, but signaling mechanisms involved are poorly understood. Data here demonstrate that flow stimulates signaling mechanisms in endothelial cells that are similarly dependent on PC-1 and PC-2. When combined with the results of our earlier study, it is evident that flow-mediated signaling and vasodilation are similarly attenuated in endothelial cells and arteries of *Pkd1* ecKO, *Pkd2* ecKO and *Pkd1/Pkd2* ecKO mice, further strengthening this conclusion (6). These results include that inducible, endothelial cell-specific knockout of either PC-1 or PC-2 similarly inhibits flow-mediated: 1) biphasic currents in endothelial cells, 2) Ca^2+^ influx-dependent cation currents in endothelial cells, 3) arterial hyperpolarization, 4) vasodilation over a broad shear stress range and 5) vasodilation that occurs via NOS and SK channel activation. We have previously demonstrated that flow stimulates PC-2-dependent Ca^2+^ influx that activates apamin-sensitive currents in endothelial cells and eNOS serine1176 phosphorylation in mesenteric arteries, consistent with our data obtained here in *Pkd1* ecKO mice (6). In contrast, PC-1 and PC-2 do not contribute to ACh-induced vasodilation, indicating that their knockout does not cause generalized endothelial cell dysfunction. Myogenic tone, depolarization-induced constriction and passive diameter are similar in *Pkd1* ecKO, *Pkd2* ecKO, *Pkd1/Pkd2* ecKO and control mouse arteries, illustrating that smooth muscle function is also not altered in these knockout mouse models.

Virtually all vertebrate cells possess at least one primary cilia, a tiny (~0.2-0.5 μm width, 1-12 μm length) immotile organelle electrically distinct from the cell body (37–39). Cilia generate compartmentalized signaling and maintain an intracellular Ca^2+^ concentration distinct from the cell soma (37–39). A great deal of debate has taken place over whether cells sense external fluid flow through proteins located in primary cilia or the plasma membrane. Primary cilia act as flow sensors in the embryonic node and in kidney collecting duct cells (40, 41). In contrast, flow does not activate Ca^2+^ influx in primary cilia of cultured kidney epithelial cells, kidney thick ascending tubules, embryonic node crown cells or in the kinocilia of inner ear hair cells (42). Instead, flow stimulates a cytoplasmic Ca^2+^ signal in the cell body that propagates to cilia to elevate intraciliary Ca^2+^ concentration (42). Here, flow stimulated plasma membrane cation currents, indicating that flow-sensing takes place in the cell body of endothelial cells.

Physiological functions of PC-1 and the potential involvement of PC-2 in PC-1-mediated signaling mechanisms in endothelial cells were unclear. Whether physical coupling of PC-1 and PC-2 is required for these proteins to traffic to the surface and to activate cellular signaling has also been a matter of debate. Studies that aimed to answer this question primarily used renal epithelial cells or recombinant proteins expressed in cultured cells or *Xenopus* oocytes. Ciliary localization of PC-2 is necessary for flow-sensing in perinodal crown cells and left-right symmetry in mouse embryos (40). In contrast, other studies demonstrated that a physical interaction between PC-1 and PC-2 is essential for these proteins to traffic to the cell surface and generate plasma membrane currents (25, 29–31,43–45). It was essential to include an IgX-chain secretion sequence into recombinant PC-1 to induce its trafficking and associated PC-2 to the plasma membrane in HEK293 cells (45). We show that when both PC-1 and PC-2 are expressed, they readily traffic to the plasma membrane in endothelial cells. PC-1 and PC-2 protein clusters are present in the endothelial cell plasma membrane and approximately one-third overlap, supporting their coassembly. Knockout of either PC-1 or PC-2 does not alter the total amount of the other polycystin protein but abolishes surface expression of both proteins. Whether PC-1 and PC-2 traffic to the plasma membrane as a complex or traffic independently and then assemble at the surface is not clear. Flow did not alter the properties of surface PC-1 or PC-2 clusters, or their intercluster distance or overlap, suggesting that flow activates a heteromeric PC-1/PC-2 complex in the plasma membrane. A previous study showed that PC-2 immunolabeling colocalized with α-tubulin, a ciliary marker, in endothelial cells of embryonic (E15.5) conduit vessels (46). We did not examine if PC-1 or PC-2 are present in cilia or if flow activates PC-1- or PC-2-dependent currents in the cilia of endothelial cells. Future studies should investigate these possibilities.

PC-1 and PC-2 did not generate currents in the plasma membrane of renal collecting duct cells or following heterologous expression in cell lines (14, 15, 17, 30, 31, 39). Knockout of either PC-1 or PC-2 also did not alter plasma membrane currents in inner medullary collecting duct (pIMCD) epithelial cells (14). Recently, it was demonstrated that plasma membrane PC-1/PC-2 channels are silent but can be measured when a gain-of-function mutation (F604P) is introduced into PC-2 (45, 47). We show that in the absence of flow, surface PC-1/PC-2 channels generate little current in endothelial cells (6). Rather, flow-activates plasma membrane currents in endothelial cells that are similarly attenuated by either PC-1 or PC-2 knockout or by the introduction of peptides that correspond to the C-terminus coiled-coil domains that interact on each protein (here and ref. (6)). Flow-mediated intracellular Ca^2+^ elevations and NO production were attenuated in cultured embryonic aortic endothelial cells from *Pkd1* and Tg737^orpk/orpk^ global knockout mice (48). As Tg737^orpk/orpk^ global knockout mice have shorter cilia or no cilia, the authors proposed that PC-1 located in cilia is a flow sensor in embryonic endothelial cells (48). In contrast, more recent studies have demonstrated that flow did not elicit Ca^2+^ signaling in primary cilia and homomeric PC-2 channels did not require PC-1 in primary cilia of pIMCD epithelial cells (14). Our data support the conclusion that flow activates PC-1/PC-2-dependent cation currents in the plasma membrane of endothelial cells.

Multiple different domains in PC-1 and PC-2 physically interact. Several groups have demonstrated that PC-1 and PC-2 couple via their C-terminal coiled-coils (26, 28, 29, 32). Recombinant PC-1 and PC-2 lacking coiled-coils can also interact via N-terminal loops (49, 50). The structure of a PC-1/PC-2 heterotetramer resolved using cryo-EM indicated that a region between TM6 and TM11 of PC-1 interdigitates with PC-2 (25). PC-1 is proposed to act both as a dominant-negative subunit in PC-1/PC-2 channels and to increase Ca^2+^ permeability over that in PC-2 homotetramers (25, 47). Our data suggest that flow stimulates physical coupling of the coiled-coil domains in PC-1/PC-2 to activate current. Alternatively, the interference peptides may uncouple constitutively bound coiled coils in PC-1/PC-2 heteromers, thereby preventing current activation by flow. It is unlikely that the interference peptides dissolve PC-1/PC-2 into individual subunits as several other domains can interact and the PC-1/PC-2 heteromeric structure was resolved in the absence of the coiled-coils (25, 50). PC-1 has been proposed to act as an atypical G protein due to the presence of a G protein-binding domain located in the C-terminus (16). The PC-1 interference peptide we used does not overlap with this G protein-binding domain, which is located between amino acids 4125 and 4143. We also show that peptides corresponding to the coiled-coil domains in either PC-1 or PC-2 similarly inhibit flow-mediated current activation. Thus, G protein signaling by PC-1 does not appear to be involved in flow-mediated PC-1/PC-2-dependent current activation in endothelial cells.

Our data suggest that flow activates Ca^2+^ influx through PC-1/PC-2 channels. Although homomeric PC-2 is a K^+^- and Na^+^-permeant channel with low Ca^2+^ permeability, heteromeric assembly of PC-2 with PC-1 increases Ca^2+^ permeability and reduces block by external Ca^2+^, supporting this concept (14, 47). We did not determine the ionic permeability of flow-activated PC-1/PC-2-dependent currents in endothelial cells (47). Endothelial cells express a wide variety of ion channels, including several other TRPs and K^+^ channels, making the isolation of a pure PC-1/PC-2-dependent current challenging. PC-1/PC-2 may also interact with other TRP channels. For example, heteromeric channels containing PC-2 and TRPM3 exhibit higher Ca^2+^ permeability than homomeric PC-2 in primary cilia of cultured renal epithelial cells (51, 52). Thus, PC-1/PC-2 channels in endothelial cells may contain other TRP subunits.

Flow may directly or indirectly activate PC-1/PC-2 in endothelial cells. PC-1 is proposed to act as a mechanical sensor and ligand-receptor in cultured kidney epithelial cells (9–11). The PC-1 extracellular N-terminus contains several putative adhesion- and ligand-binding sites that may confer mechanosensitivity (3, 7, 8). Recent evidence indicates that the C-type lectin domain located in the PC-1 N-terminus activates recombinant PC-1/PC-2 channels, providing one possible activation mechanism for flow (45). Conceivably, other mechanosensitive proteins, such as Piezo1 and GPR68, may also stimulate PC-1/PC-2-dependent currents in endothelial cells (53, 54). Investigating the mechanisms by which flow activates PC-1/PC-2-dependent currents should be a focus of future studies.

ADPKD is typically characterized by the appearance of renal cysts, but patients can develop hypertension prior to any kidney dysfunction, with cardiovascular disease the leading (~50 %) cause of death in patients (20–24). Our study raises the possibility that hypertension and cardiovascular disease in ADPKD patients may involve altered PC-1/PC-2 function in endothelial cells. Consistent with our data, human ADPKD patients exhibit loss of nitric oxide release and a reduction in endothelium-dependent dilation during increased blood flow (6, 55). Hypertension in humans is also associated with endothelial dysfunction and attenuated flow-mediated dilation (56). Conceivably, hypertension may also be associated with dysfunctional PC-1/PC-2 signaling in endothelial cells. Our demonstration that PC-1/PC-2 elicits vasodilation may lead to studies identifying potential dysfunction in patients with hypertension, ADPKD and other cardiovascular diseases.

In summary, we show that PC-1 regulates arterial contractility through the formation of an interdependent plasma membrane signaling complex with PC-2 in endothelial cells. Flow stimulates PC-1/PC-2-dependent currents in endothelial cells, leading to Ca^2+^ influx, NOS and SK channel activation, vasodilation and a reduction in blood pressure.

## Materials and Methods

### Animals

All animal studies were performed in accordance with the Institutional Animal Care and Use Committee (IACUC) at the University of Tennessee Health Science Center. *Pkd1^fl/fl^* and *Pkd2^fl/fl^* mice were obtained from the Baltimore PKD Center (Baltimore, MD). *Cdh5*(PAC)-creERT2 mice were a kind gift from Cancer Research UK (57). *Pkd1^fl/fl^* mice were crossed with tamoxifen-inducible endothelial cell-specific Cre mice (Cdh5(PAC)-CreERT2, Cancer Research UK) to generate *Pkd1^fl/fl^:Cdh5(PAC)-CreERT2* mice. *Pkd2^fl/fl^:Cdh5(PAC)-CreERT2* mice were generated as previously described (6). *Pkd1^fl/fl^* mice were crossed with *Pkd2^fl/fl^* mice to produce *Pkd1^fl/fl^/Pkd2^fl/fl^* mice. *Pkd1^fl/fl^:Cdh5(PAC)-CreERT2* mice were crossed with *Pkd2^fl/fl^:Cdh5(PAC)-CreERT2* mice to generate *Pkd1^fl/fl^/Pkd2 ^fl/fl^:Cdh5(PAC)-CreERT2* mice. The genotypes of all mice were confirmed using PCR (Transnetyx, Memphis, TN) before use. *Pkd1^fl/fl^, Pkd2^fl/fl^* and *Pkd1^fl/fl^/Pkd2 ^fl/fl^* cre negative mice were used as controls. All mice (male, 12 weeks of age) were injected with tamoxifen (50 mg/kg, i.p.) once per day for 5 days and studied between 7 and 14 days after the last injection.

### Tissue preparation and endothelial cell isolation

Mesenteric artery branches from first to fifth order were cleaned of adventitial tissue and placed into ice-cold physiological saline solution (PSS) that contained (in mM): 112 NaCl, 6 KCl, 24 NaHCO_3_, 1.8 CaCl_2_, 1.2 MgSO_4_, 1.2 KH_2_PO_4_ and 10 glucose, gassed with 21% O_2_, 5% CO_2_ and 74% N_2_ (pH 7.4). Endothelial cells were dissociated by introducing endothelial cell basal media (Endothelial cell GM MV2, Promocell) containing 2 mg/ml collagenase type 1 (Worthington Biochemical) into the arterial lumen and left to incubate for 30-40 minutes at 37°C. Endothelial cells were either studied immediately (fresh-isolated) or cell isolate was placed into endothelial cell basal media containing growth supplements (Promocell) that support endothelial cell survival. Primary-cultured endothelial cells were studied within 5 days of isolation.

### Genomic PCR

Genomic DNA was isolated from mesenteric arteries using a Purelink Genomic DNA kit (Thermo Fisher Scientific). PCR was then performed on an Eppendorf Gradient thermal cycler using the following protocol: 95°C for 2 min, then 35 cycles of 95°C for 30 s, 56°C for 30 s, and 72°C for 30 s. Primer sequences were as follows: *Pkd1^fl/fl^* GTTATTCGAGGTCGCTAGACCCTATC (forward), GTTACAGATGAGGCCCAGGGAAAG (reverse), *Pkd1* ecKO: GGTACGAGAGAGAAGTGGTCTCAGGA (forward), GAGATCCCACCGCGGTTTTGCTAGAAGGCA (reverse). PCR products were separated on 1.5% agarose gels.

### Western Blotting

Mesenteric artery segments comprising second-to fifth-order vessels were used for Western blotting. Arteries were transferred to an eppendorf tube containing RIPA Buffer (Sigma: R0278) and protease inhibitor cocktail (Sigma: P8340, 1 in 100 dilution). Arteries were cut into small segments using micro scissors and mechanically broken down using a homogenizer (Argos Technologies: A0001). The lysate was centrifuged at 4°C, 10,000 rpm for 2 minutes. This process was repeated 3 times, after which the supernatant was collected. Proteins in lysate were separated on 7.5% SDS-polyacrylamide gels and blotted onto nitrocellulose membranes. Membranes were blocked with 5% milk and incubated with one of the following primary antibodies: PC-1 (Santa Cruz), PC-2 (Santa Cruz), Piezo1 (Proteintech), GPR68 (NOVUS), SK3 (Sigma (P0608)), IK (Alomone), TRPV4 (MilliporeSigma), eNOS (Abcam), or actin (MilliporeSigma) overnight at 4°C. Membranes were washed and incubated with horseradish peroxidase-conjugated secondary antibodies at room temperature. Protein bands were imaged using a ChemiDoc Touch Imaging System (Bio-Rad), quantified using ImageJ software and normalized to actin.

### Laser-scanning confocal microscopy

Arteries were cut open longitudinally and fixed with 4% paraformaldehyde in PBS for 1 hr. Following a wash in PBS, arteries were permeabilized with 0.2% Triton X-100 and blocked with 3% BSA + 5% serum for 1 hour at room temperature. For en face imaging experiments, arteries were incubated overnight with anti-PC-1 monoclonal primary antibody (E3 5F4A2, Baltimore PKD Center) and anti-CD31 primary monoclonal antibody (Abcam 7388) at 4°C. Arteries were then incubated with Alexa Fluor 488 donkey anti-rat, Alex Fluor 546 donkey anti-mouse secondary antibodies (1:500; Molecular Probes) and 4’,6-diamidino-2-phenylindole, dihydrochloride (DAPI) (1:1000; Thermo Scientific) for 1 hour at room temperature. After washing with PBS, arteries were mounted in 80% glycerol solution. DAPI, Alexa 488 and Alexa 546 were excited at 405, 488 and 561 nm with emission collected at ≤460 nm and ≥500 nm, respectively, using a Zeiss LSM 710 laser-scanning confocal microscope.

### Patch-clamp electrophysiology

The conventional whole-cell configuration was used to measure steady-state currents in primary-cultured endothelial cells at a holding potential of −60 mV. For experiments using physiological ionic gradients, the bath solution contained (in mM): NaCl 134, KCl 6, HEPES 10, MgCl_2_ 1, CaCl_2_ 2 and glucose 10 (pH 7.4). The Ca^2+^-free bath solution was the same composition as bath solution except Ca^2+^ was omitted and 1 mM EGTA was included. The pipette solution contained (in mM): K aspartate 110, KCl 30, HEPES 10, glucose 10, EGTA 1, Mg-ATP 1 and Na-GTP 0.2, with total MgCl_2_ and CaCl_2_ adjusted to give free concentrations of 1 mM Mg^2+^ and 200 nM Ca^2+^, respectively (pH 7.2). For experiments measuring I_Cat_, the bath solution contained (in mM): Na aspartate 135, NaCl 5, HEPES 10, glucose 10 and MgCl_2_ 1 (pH 7.4). A low Na^+^ bath solution used when measuring I_Cat_ was (in mM): NMDG-asp 135, NaCl 5, HEPES 10, glucose 10 and MgCl_2_ 1 (pH 7.4). The pipette solution contained (in mM): Na aspartate 135, NaCl 5, HEPES 10, glucose 10, EGTA 1, Mg-ATP 1 and Na-GTP 0.2, with total Mg^2+^ and Ca^2+^ adjusted to give free concentrations of 1 mM and 200 nM, respectively (pH 7.2). PC-1 and PC-2 coiled-coil domain peptides were custom-made (Genscript). PC-1 or PC-2 peptides were added to pipette solution immediately before use at a final concentration of 1 μM. The peptide sequence corresponding to PC-1 is FDRLNQATEDVYQLEQQL and for PC-2 is KRREVLGRLL. Free Mg^2+^ and Ca^2+^ were calculated using WebmaxC Standard (https://somapp.ucdmc.ucdavis.edu/pharmacology/bers/maxchelator/webmaxc/webmaxcS.htm). The osmolarity of solutions was measured using a Wescor 5500 Vapor Pressure Osmometer (Logan, UT, USA). Currents were filtered at 1 kHz and digitized at 5 kHz using an Axopatch 200B amplifier and Clampex 10.4 (Molecular Devices. Offline analysis was performed using Clampfit 10.4. The flow-activated transient inward current was measured at its peak in each cell. The steady-state inward current was the average of at least 45 seconds of contiguous data.

### Pressurized artery membrane potential measurements

Membrane potential was measured by inserting sharp glass microelectrodes (50-90 MΩ) filled with 3 M KCl into the adventitial side of pressurized third- and fourth-order mesenteric arteries. Membrane potential was recorded using a WPI FD223a amplifier and digitized using a MiniDigi 1A USB interface, pClamp 9.2 software (Axon Instruments) and a personal computer. Criteria for successful intracellular impalements were a sharp negative deflection in potential on insertion, stable voltage for at least 1 min after entry, a sharp positive voltage deflection on exit from the recorded cell and a <10% change in tip resistance after impalement.

### Pressurized artery myography

Experiments were performed using isolated third- and fourth-order mesenteric arteries using PSS gassed with 21% O_2_/5% CO_2_/74% N_2_ (pH 7.4). Arterial segments 1-2 mm in length were cannulated at each end in a perfusion chamber (Living Systems Instrumentation) continuously perfused with PSS and maintained at 37°C. Intravascular pressure was altered using a Servo pump model PS-200-P (Living systems) and monitored using pressure transducers. Following development of stable myogenic tone, luminal flow was introduced during experiments using a P720 peristaltic pump (Instech). Arterial diameter was measured at 1 Hz using a CCD camera attached to a Nikon TS100-F microscope and the automatic edge-detection function of IonWizard software (Ionoptix). Myogenic tone was calculated as: 100 x (1-D_active_/D_passive_) where D_active_ is active arterial diameter and D_passive_ is the diameter determined in the presence of Ca^2+^-free PSS supplemented with 5 mM EGTA.

### Telemetric blood pressure and locomotion measurements

Telemetric blood pressure recordings were performed at the University of Tennessee Health Science Center. Briefly, transmitters (PA-C10, Data Sciences International) were implanted subcutaneously into anesthetized mice, with the sensing electrode placed in the aorta via the left carotid artery. Blood pressure was recorded every 20 s for 5 days prior to tamoxifen injection and every 20 s for the entire time period between 7 and 29 days after the last tamoxifen injection (50 mg/kg per day for 5 consecutive days, i.p) using a PhysioTel Digital telemetry platform (Data Sciences International). Dataquest A.R.T. software was used to acquire and analyze data.

### Kidney histology

Kidney sections were stained with H&E and examined by Probetex, Inc (San Antonio, Texas). Briefly, image analysis was performed to measure the glomerular size and tubular crosssectional diameter. The glomerular size was measured by tracing the circumference of 25 random glomeruli and surface area was calculated using the polygonal area tool of Image-Pro 4.5 image analysis software calibrated to a stage micrometer. Tubular size was measured using the linear length tool of Image-Pro 4.5 imaging software. The tracing tool was applied at the diameter of cross-sectional profiles of 5 proximal tubules/image (total of 25/section). Glomerular and tubular images were calibrated to a stage micrometer and data was transferred to an Excel spreadsheet and statistical analysis was performed by Excel analysis pack.

### Co-immunoprecipitation

Mesenteric artery segments comprising second-to fifth-order vessels were used. Arteries were transferred to an eppendorf tube containing lysis buffer and protease inhibitor cocktail (Sigma: P8340, 1 in 100 dilution). Arteries were cut into small segments using micro scissors and broken down using a mechanical homogenizer (Argos Technologies: A0001). Arterial lysate was centrifuged for 2 minutes at 10,000 rpm and at 4°C. This process was repeated 3 times, after which the supernatant was collected. Proteins were pulled down from arterial lysate using a Pierce crosslink Magnetic IP/coIP kit (Thermoscientific) as per the manufacturer’s instructions. Samples were incubated with PC-2 antibody (1:20, Santa Cruz) that was covalently bound to protein A/G Magnetic Beads (Pierce) overnight at 4 °C. Following washing and elution, immunoprecipitates were analyzed using Western blotting.

### Immunofluorescence Resonance Energy Transfer (ImmunoFRET) imaging

Primary-cultured mesenteric artery endothelial cells were fixed with paraformaldehyde and permeabilized with 0.1% Triton X-100 for 2 min at room temperature. After blocking with 5% bovine serum albumin (BSA), the cells were treated overnight at 4°C with anti PC-1 (Rabbit polyclonal, Santa Cruz, sc-25570, 1:100 dilution, RRID:AB_2163357) and anti PC-2 (mouse monoclonal, Santa Cruz, sc-28331, 1:100 dilution, RRID:AB_672377) antibodies in PBS containing 5% BSA. After a wash, cells were incubated for 1 h at 37°C with secondary antibodies: Goat anti-Mouse Alexa Fluor 488 (ThermoFisher, A-11001) and Donkey anti-Rabbit Alexa Fluor 555 (ThermoFisher, A-31572). Coverslips were then washed and mounted on glass slides. Fluorescence images were acquired using a Zeiss 710 laser-scanning confocal microscope. Alexa 488 and Alexa 546 were excited at 488 and 543 nm and emission collected at 505-530 and >560 nm, respectively. Images were acquired using a z-resolution of ~1 μm. Images were background-subtracted and normalized FRET (N-FRET) was calculated on a pixel-by-pixel basis for the entire image and in regions of interest (within the boundaries of the cell) using the Xia method (41) and Zeiss LSM FRET Macro tool version 2.5 as previously described (48).

### Lattice Structured Illumination Microscopy (Lattice-SIM)

Arteries were cut open longitudinally and fixed with 4% paraformaldehyde in PBS for 1 hr. Following a wash in PBS, arteries were permeabilized with 0.2% Triton X-100 and blocked with 3% BSA + 5% serum for 1 hour at room temperature. Arteries were incubated overnight with anti-PC-1 monoclonal primary antibody (E3 5F4A2, Baltimore PKD Center), anti-PC-2 monoclonal antibody (3374 CT-1 414, Baltimore PKD Center) and anti-CD31 primary monoclonal antibody (Abcam 7388) at 4°C. Arteries were then incubated with Alexa Fluor 488 donkey anti-mouse, Alex Fluor 546 donkey anti-rabbit secondary antibodies and Alex Fluor 647 goat anti-rat secondary antibodies (1:500; Molecular Probes) for 1 hour at room temperature. After washing with PBS, arteries were mounted in 80% glycerol solution. Lattice-SIM images were acquired on a Zeiss Elyra 7 AxioObserver microscope equipped with 63x PlanApochromat (NA 1.46) oil immersion lens and an sCMOS camera. Lattice-SIM reconstruction was performed using the SIM processing Tool of Zeiss ZEN Black 3.0 SR software. Colocalization analysis of Lattice-SIM data was performed using Pearson’s and Mander’s coefficients.

### Single-Molecule Localization Microscopy (SMLM)

Primary-cultured mesenteric artery endothelial cells were seeded onto 35 mm glass-bottom dishes (MaTeK Corp). Cells were fixed in 4 % paraformaldehyde, permeabilized with 0.5 % Triton X-100 phosphate-buffered saline (PBS) solution and then blocked with 5 % BSA. Cells were immunolabelled using primary antibodies to PC-1 (Baltimore PKD Center), PC-2 (Baltimore PKD Center) and CD31 (Abcam 7388) overnight at 4°C. Alexa Fluor 488 donkey antimouse, Alexa Fluor 555 goat anti-rat and Alexa Fluor 647 donkey anti-rabbit secondary antibodies were used for detection. An oxygen scavenging, thiol-based photo-switching imaging buffer was used (GLOX: 50% glucose, 10% PBS, 24 mg/ml glucose oxidase, 12.6 mg/ml catalase supplemented with a reducing agent (cysteamine hydrochloride-MEA) at 100 mM, pH 7.8). Cells were imaged using a super-resolution Zeiss Elyra 7 microscope using 488 nm (500mW), 561 nm (500mW) and 642 nm (500mW) lasers. A 63x Plan-Apochromat (NA 1.46) oil immersion lens and a CMOS camera were used to acquire images. The camera was run in frame-transfer mode at a rate of 100 Hz (30 ms exposure time). Fluorescence was detected using TIRF mode with emission band-pass filters of 550-650 and 660-760 nm.

Localization precision was calculated using σ^2^_x,y_ = [(s^2^ + q^2^/12) / N] + [(8πs^4^b^2^) / (q^2^N^2^)](1) σ^2^_x,y_ ≈ s^2^ / N(2), where σ_x,y_ is the localization precision of a fluorescent probe in lateral dimensions, s is the standard deviation of the point-spread function, N is the total number of photons gathered, q is the pixel size in the image space, and b is the background noise per pixel. The precision of localization is proportional to DLR/√N, where DLR is the diffraction-limited resolution of a fluorophore and N is the average number of detected photons per switching event, assuming the point-spread functions are Gaussian. PC-1 and PC-2 localization were reconstructed from 35,000-40,000 images based on fitting signals to a Gaussian function and taking into account a point spread function calculated using a standardized 40 nm bead slide (ZEN Black software, Zeiss). The first 5000 frames were excluded from the reconstruction to account for time to stabilize photo-switching of the probes. Software drift correction was applied using a model-based cross-correlation.

### Statistical analysis

OriginLab and GraphPad InStat software were used for statistical analyses. Values are expressed as mean ± SEM. Student t-test was used for comparing paired and unpaired data from two populations and ANOVA with Holm-Sidak post hoc test used for multiple group comparisons. P<0.05 was considered significant. Power analysis was performed to verify that the sample size gave a value > 0.8 if P was > 0.05. Kidney histology and blood pressure experiments were all done single-blind, wherein the person performing the experiments and analysis of the results was not aware of the mouse genotype.

## Acknowledgments

This work was supported by NIH/NHLBI grants HL133256, HL137745, HL155180 and HL155186 (to J.H.J), HL19134-46 (to K.U.M) and an American Heart Association (AHA) Postdoctoral Fellowship (20POST35210200) and Career Development Award R073037556 (to C.M.). We thank Dr. Simon Bulley for initial breeding of mouse lines and Dr. Manuel Navedo (UC, Davis) for help with Image J software.

## Supplementary Information

**Fig. S1.**
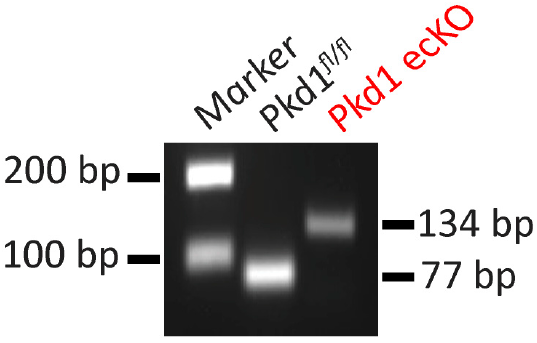
Genomic PCR indicating that tamoxifen stimulated Cre-recombination in mesenteric arteries of *Pkd1^fl/fl^: Cdh5*(PAC)-creERT2 mice. Representative of four separate experiments.

**Fig. S2.**
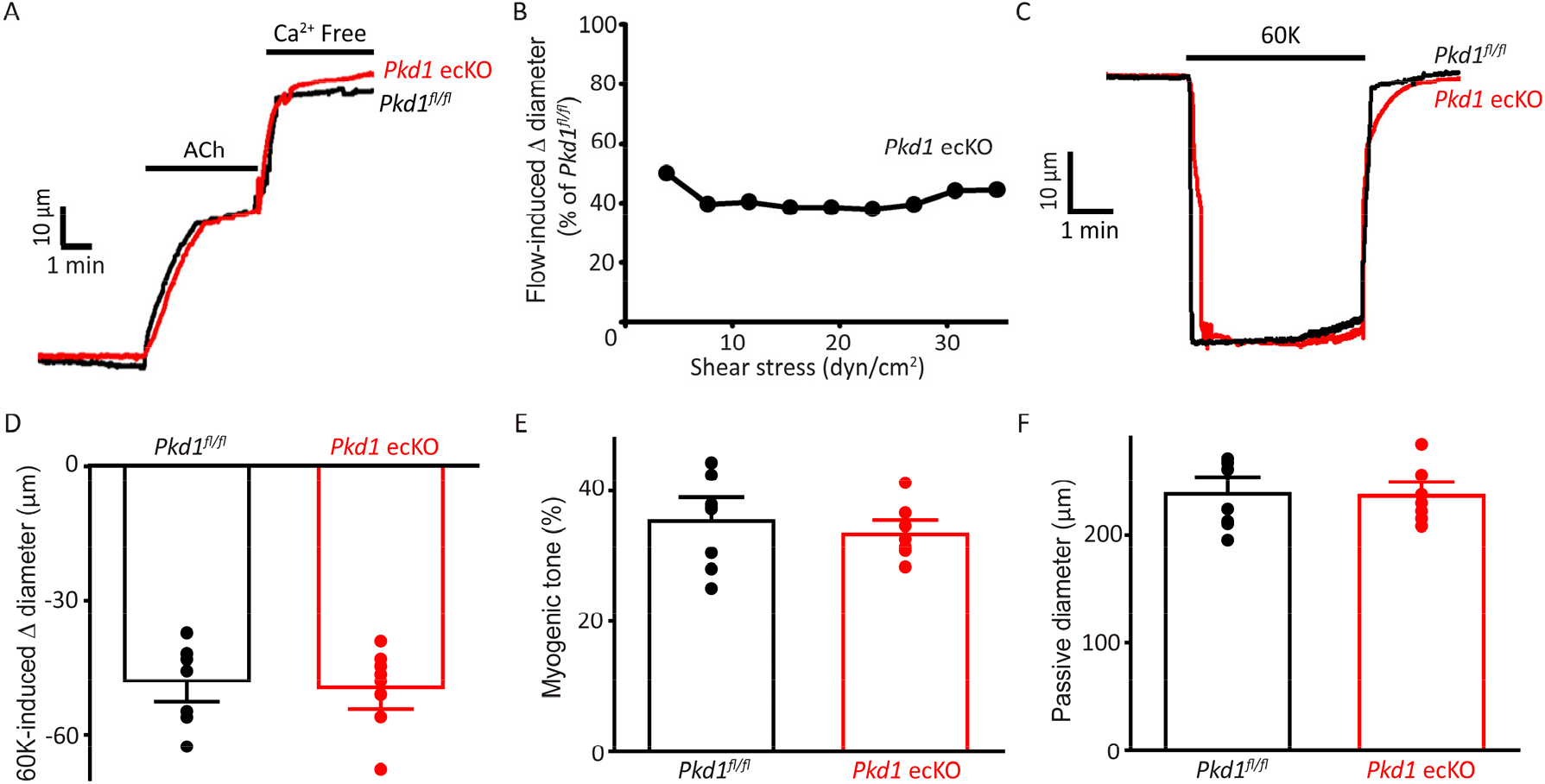
Depolarization-induced vasoconstriction and passive diameter are similar in *Pkd1^fl/fl^* and *Pkd1* ecKO arteries. (A) Representative traces illustrating dilations to ACh (10 μM) and Ca^2+^-free bath solution in pressurized (80 mmHg) *Pkd1^fl/fl^* and *Pkd1* ecKO arteries. (B) PC-1 knockout attenuates flow-mediated dilation over a broad shear stress range in pressurized (80 mmHg) mesenteric arteries. n=4 for each shear stress value. (C) Representative traces illustrating constriction to 60 mM K^+^ in pressurized (10 mmHg) arteries. (D) Mean data for 60 mM K^+^-induced constriction. n=8 for each dataset. (E) Mean myogenic tone at 80 mmHg. n=8 for each dataset. (F) Mean data for passive diameter (Ca^2+^-free PSS) in pressurized (80 mmHg) arteries. n=8 for each.

**Fig. S3.**
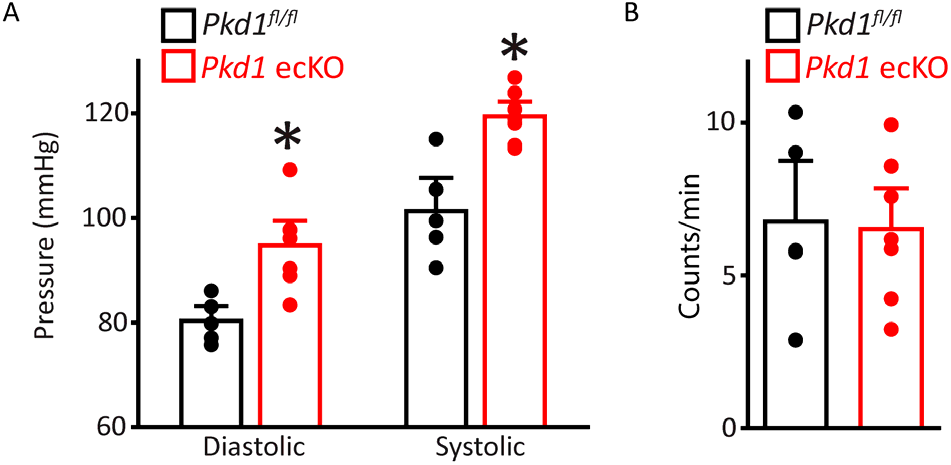
Diastolic and systolic blood pressures are higher in *Pkd1* ecKO mice. (A) Mean diastolic and systolic blood pressures for *Pkd1^fl/fl^* (n=5) and *Pkd1* ecKO (n=7) mice. *=P<0.05 vs *Pkd1^fl/fl^.* (B) Mean data for *Pkd1^fl/fl^* (n=5) and *Pkd1* ecKO (n=7) mice activity.

**Fig. S4.**
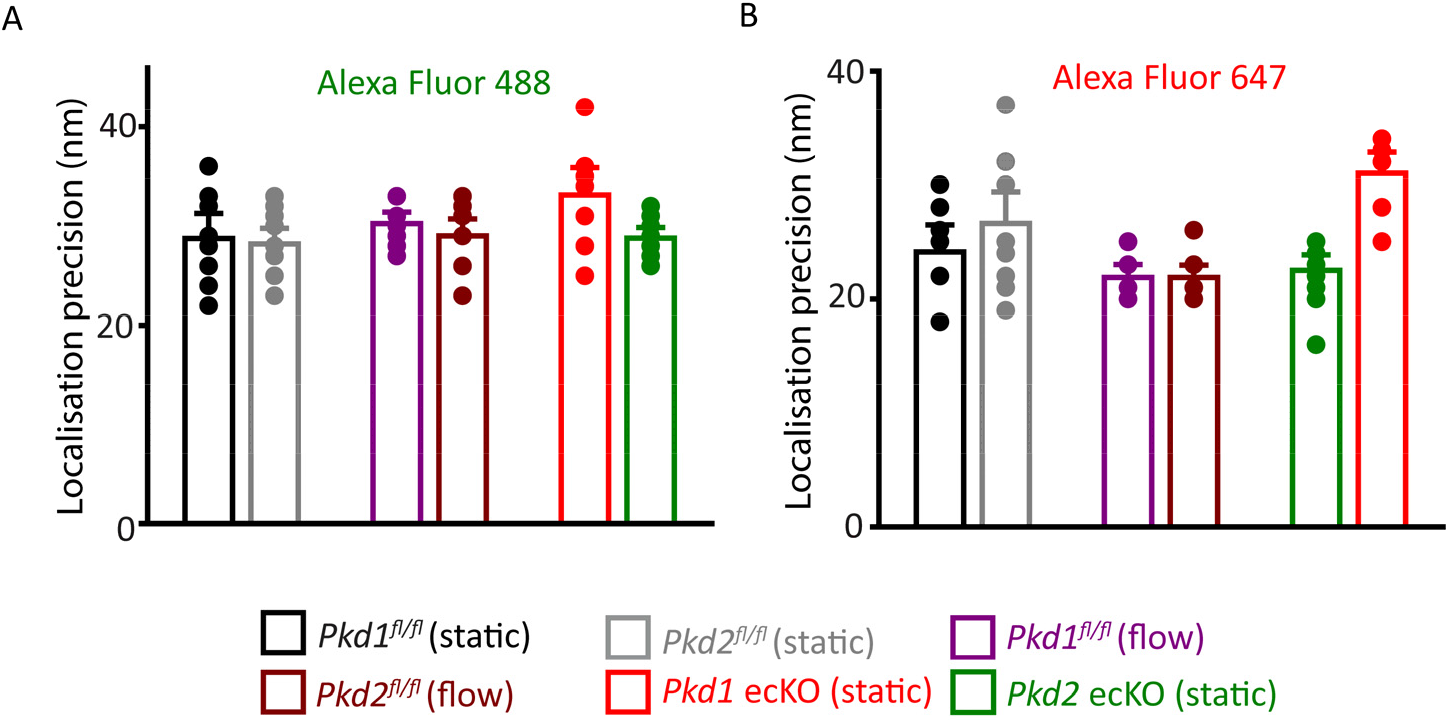
Localization precision of fluorophores used in SMLM experiments. Mean data illustrating the localization precision (FWHM) of Alexa Fluor 488 (panel A)- and Alexa Fluor 647 (panel B)-tagged secondary antibodies in endothelial cells under all conditions when imaged using SMLM. (A) Experimental numbers for Alexa Fluor 488 in *Pkd1^fl/fl^* (static), *Pkd2^fl/fl^* (static), *Pkd1^fl/fl^* (flow), *Pkd2^fl/fl^* (flow), *Pkd1* ecKO (static), *Pkd2* ecKO (static) are 8, 10, 8, 8, 8 and 9, respectively. (B) Experimental numbers for Alexa Fluor 647 in *Pkd1^fl/fl^* (static), *Pkd2^fl/fl^* (static), *Pkd1^fl/fl^* (flow), *Pkd2^fl/fl^* (flow), *Pkd1* ecKO (static), *Pkd2* ecKO (static) are 8, 10, 8, 8, 9 and 8, respectively. Numbers provided in the results are the combined means of all data collected for Alexa Fluor 488 and Alexa Fluor 647 in all genotypes in static and flow.

**Fig. S5.**
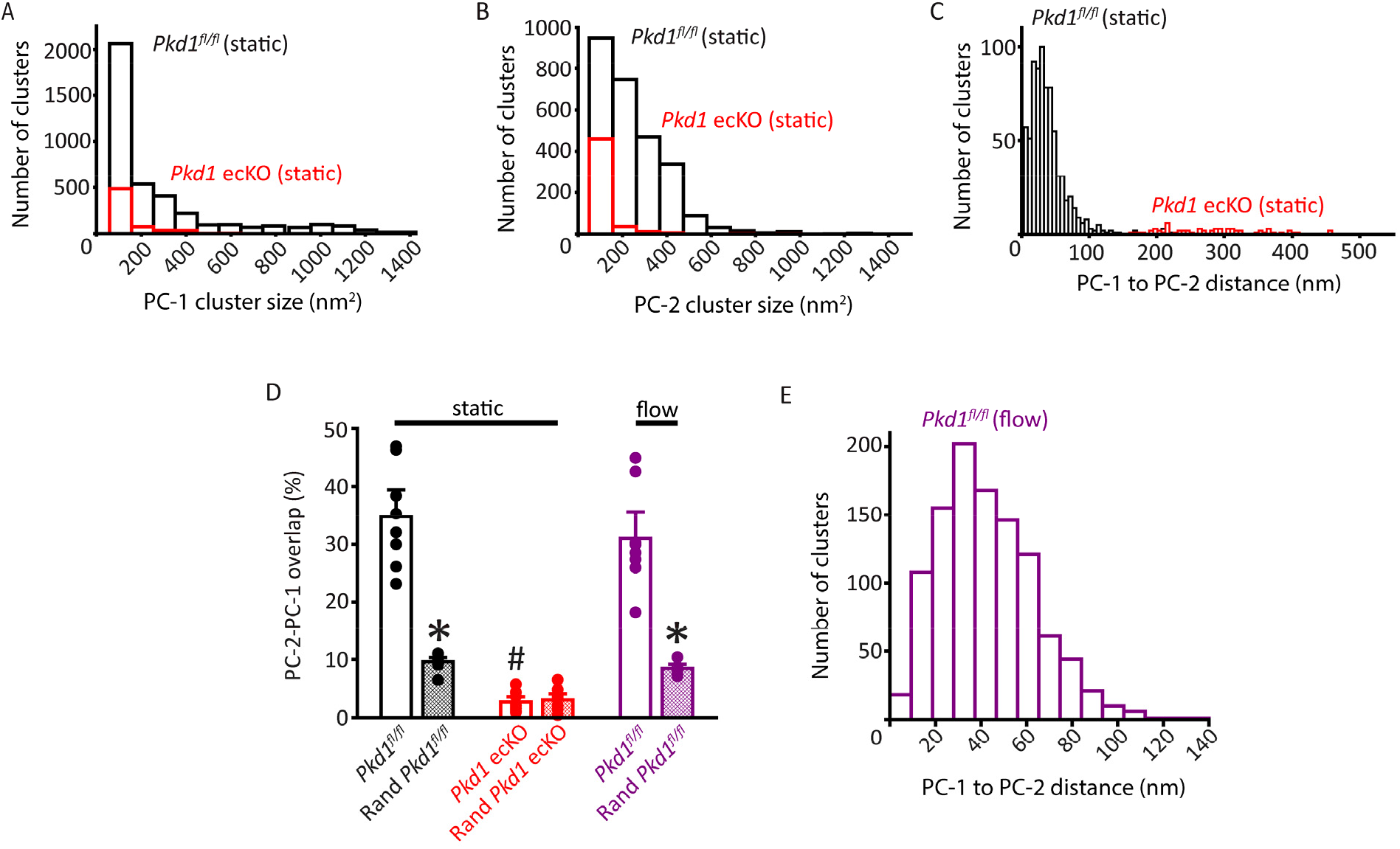
Properties of PC-1 and PC-2 clusters and their spatial proximity and overlap in *Pkd1^fl/fl^* and *Pkd1* ecKO endothelial cells. (A) Histogram illustrating PC-1 cluster size in a *Pkd1^fl/fl^* (static) and *Pkd1* ecKO (static) endothelial cell. (B) Histogram illustrating PC-2 cluster size in a *Pkd1^fl/fl^* (static) and *Pkd1* ecKO (static) endothelial cell. (C) Histogram illustrating the distance from each PC-1 cluster to the nearest PC-2 neighbor in a *Pkd1^fl/fl^* and *Pkd1* ecKO endothelial cell. (D) Mean experimental and randomized data for PC-2 to PC-1 overlap in *Pkd1^fl/fl^* cells in static and flow and *Pkd1* ecKO cells in static. n=8 for each dataset. * P<0.05 versus respective floxed control in static condition, # P<0.05 versus *Pkd1*^fl/f^ static. (E) Histogram illustrating the distance from each PC-1 cluster to the nearest PC-2 neighbor in a *Pkd1^fl/fl^* endothelial cell under flow (10 ml/min).

**Fig. S6.**
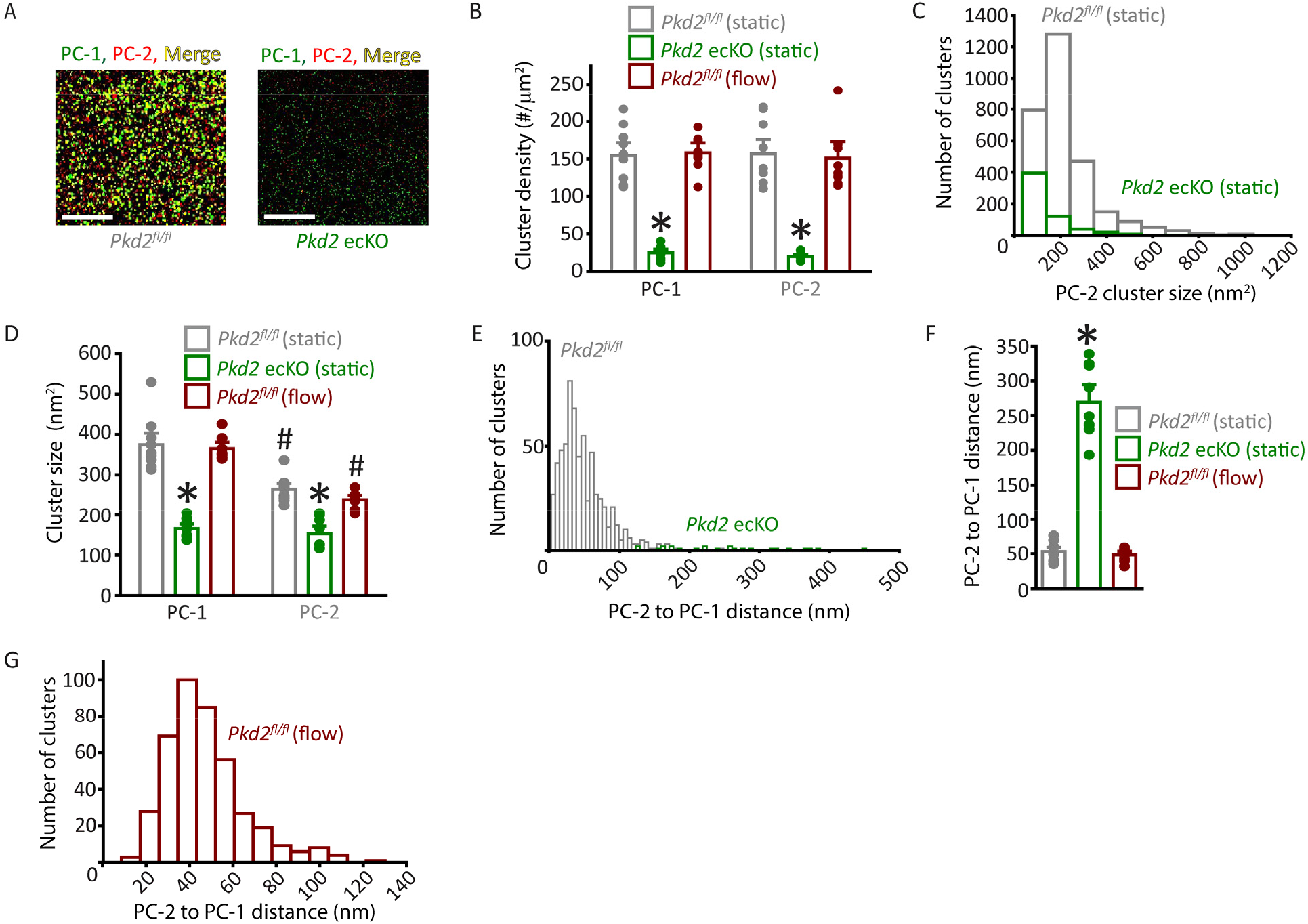
Properties of PC-1 and PC-2 surface clusters in endothelial cells of *Pkd2^fl/fl^* and *Pkd2* ecKO mice. (A) TIRF-SMLM images of PC-1 and PC-2 surface clusters in a *Pkd2^fl/fl^* and *Pkd2* ecKO endothelial cell. Scale bars = 5 μm. (B) Mean data for PC-1 and PC-2 cluster density measured in *Pkd2^fl/fl^* and *Pkd2* ecKO endothelial cells under static and flow conditions. Experimental numbers for *Pkd2^fl/fl^* static, *Pkd2* ecKO static and *Pkd2^fl/fl^* flow are 10, 9 and 9 for PC-1 and PC-2. * P<0.05 versus *Pkd2^fl/fl^* (static). (C) Histogram of individual PC-1 and PC-2 cluster sizes in a *Pkd2^fl/fl^* (static) and *Pkd2* ecKO (static) endothelial cell. (D) Mean data for PC-1 and PC-2 cluster sizes measured in *Pkd2^fl/fl^* and *Pkd2* ecKO endothelial cells under static and flow (10 ml/min) conditions. Experimental numbers for *Pkd2^fl/fl^* static, *Pkd2* ecKO static and *Pkd2^fl/fl^* flow are 10, 9 and 9 for PC-1 and PC-2. * P<0.05 versus same protein in *Pkd2^fl/fl^* static, # P<0.05 versus PC-1 in same condition. (E) Histogram illustrating distances from each PC-2 cluster to the nearest PC-1 neighbor in a *Pkd2^fl/fl^* and *Pkd2* ecKO endothelial cell. (F) Mean data for distance from each PC-2 cluster to its nearest PC-1 neighbor. n=10 for *Pkd2^fl/fl^* static, n=9 for *Pkd2* ecKO static, n=8 for *Pkd2^fl/fl^* flow. * P<0.05 versus *Pkd2^fl/fl^* static and *Pkd2^fl/fl^* flow. (G) Histogram illustrating the distance from each PC-2 cluster to the nearest PC-1 neighbor in a *Pkd2^fl/fl^* endothelial cell under flow (10 ml/min).

**Fig. S7.**
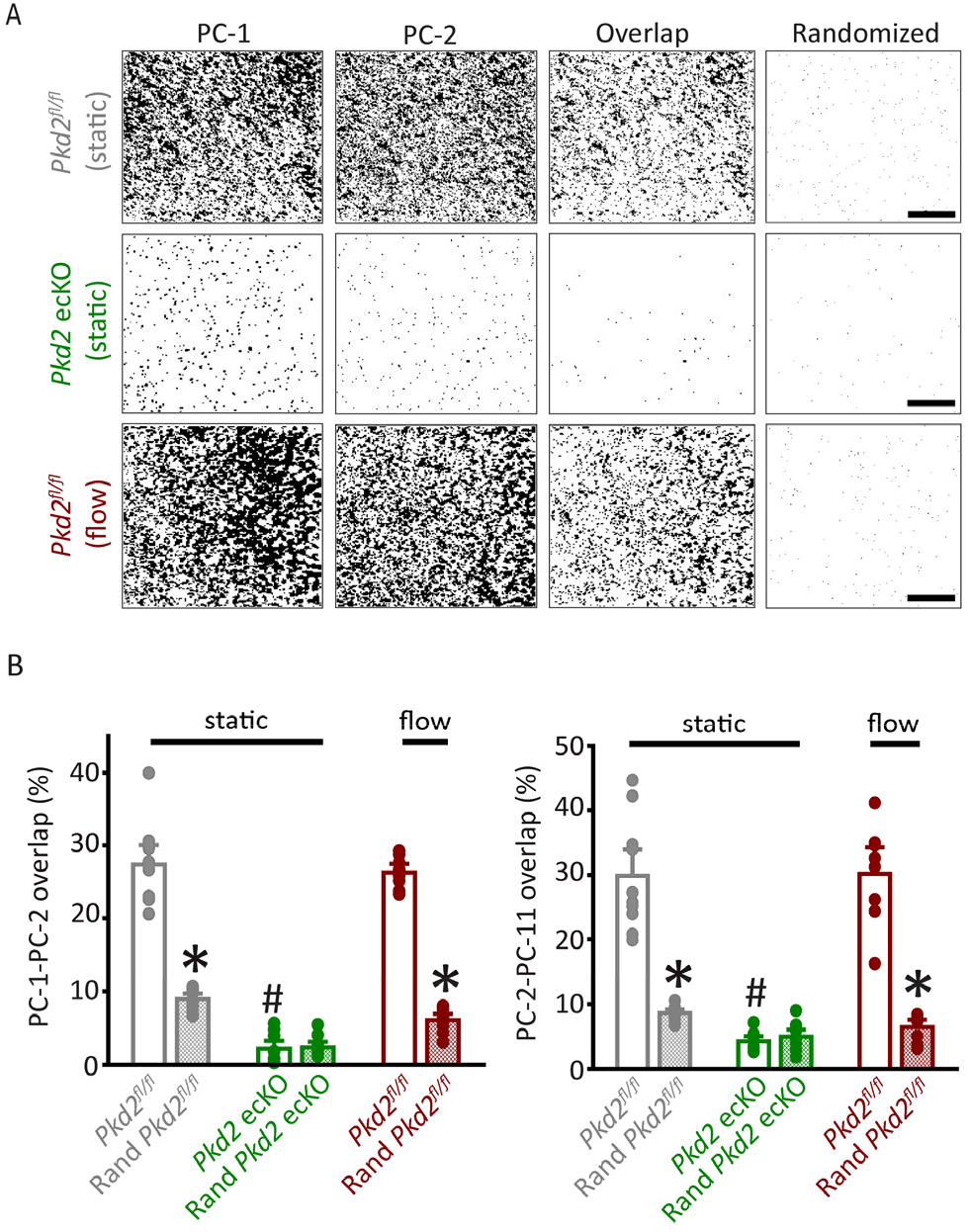
Analysis of PC-1 and PC-2 cluster overlap in endothelial cells of *Pkd2^fl/fl^* and *Pkd2* ecKO mice. (A) TIRF-SMLM images of PC-1 and PC-2 clusters, overlap of PC-1 and PC-2 data and overlap of PC-1 and PC-2 data following Costes’ randomization simulation. Scale bars = 5 μm. (B) Mean experimental and randomized overlap data for PC-1 to PC-2 and PC-2 to PC-1 in *Pkd2^fl/fl^* cells in static and flow conditions and *Pkd2* ecKO cells in static. n=10 for *Pkd2^fl/fl^* static, n=9 for *Pkd2* ecKO static, n=8 for *Pkd2^fl/fl^* flow. * P<0.05 versus respective floxed control in static condition, # P<0.05 versus *Pkd2*^fl/f^ static.

**Fig. S8.**
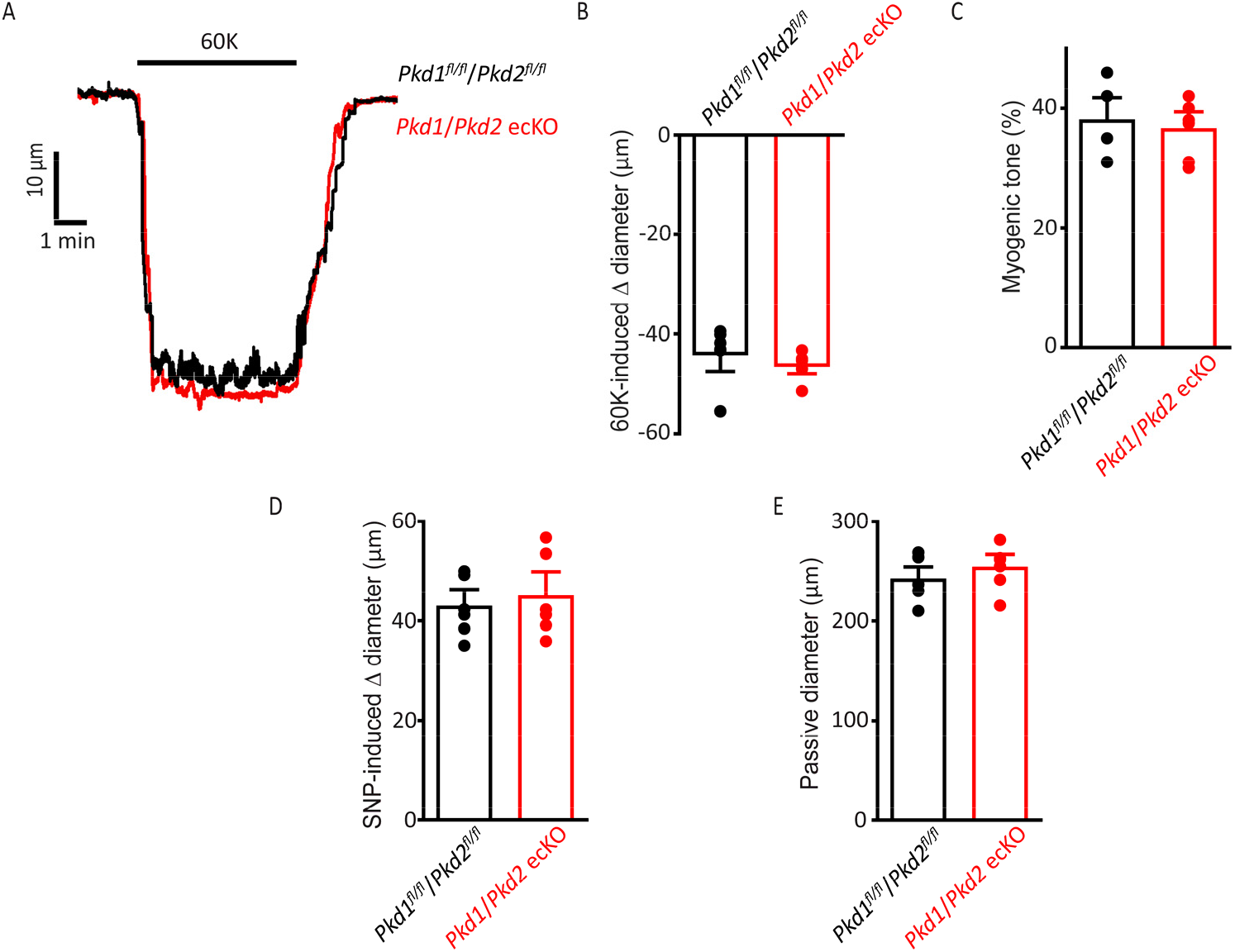
Smooth muscle-specific vasoconstriction and vasodilation and passive diameter are unaltered in *Pkd1/Pkd2* ecKO mouse arteries. (A) Representative traces illustrating 60 mM-induced K^+^ constriction in a pressurized (10 mmHg) *Pkd1^fl/fl^/Pkd2^fl/fl^* and *Pkd1/Pkd2* ecKO artery. (B) Mean data for 60 mM K^+^-induced constriction. n=6 for each dataset. (C) Mean myogenic tone in pressurized (80 mmHg) mesenteric arteries from *Pkd1^fl/fl^/Pkd2^fl/fl^* and *Pkd1/Pkd2* ecKO mice. n=6 for each dataset. (D) Mean data for SNP (10μM)-induced dilation in pressurized (10mmHg) *Pkd1^fl/fl^/Pkd2^fl/fl^* and *Pkd1/Pkd2* ecKO arteries. n=6 for each dataset. (E) Mean data for passive diameter in pressurized (80 mmHg) arteries. n=6 for each dataset.

